# A Cas9 nanoparticle system with truncated Cas9 target sequences on DNA repair templates enhances genome targeting in diverse human immune cell types

**DOI:** 10.1101/591719

**Authors:** David N. Nguyen, Theodore L. Roth, Jonathan Li, Peixin Amy Chen, Murad R. Mamedov, Linda T. Vo, Victoria Tobin, Ryan Apathy, Daniel Goodman, Eric Shifrut, Jeffrey A. Bluestone, Jennifer M. Puck, Francis C. Szoka, Alexander Marson

## Abstract

Virus-modified T cells are approved for cancer immunotherapy, but more versatile and precise genome modifications are needed for a wider range of adoptive cellular therapies^1–4^. We recently developed a non-viral CRISPR–Cas9 system for genomic site-specific integration of large DNA sequences in primary human T cells^5^. Here, we report two key improvements for efficiency and viability in an expanded variety of clinically-relevant primary cell types. We discovered that addition of truncated Cas9 target sequences (tCTS) at the ends of the homology directed repair (HDR) templates can interact with Cas9 ribonucleoproteins (RNPs) to ‘shuttle’ the template and enhance targeting efficiency. Further, stabilizing the Cas9 RNPs into nanoparticles with poly(glutamic acid) improved editing, reduced toxicity, and enabled lyophilized storage without loss of activity. Combining the tCTS HDR template modifications with polymer-stabilized nanoparticles increased gene targeting efficiency and viable cell yield across multiple genomic loci in diverse cell types. This system is an inexpensive, user-friendly delivery platform for non-viral genome reprogramming that we successfully applied in regulatory T cells (Tregs), γδ-T cells, B cells, NK cells, and primary and iPS-derived^6^ hematopoietic stem progenitor cells (HSPCs).

---

We recently reported an approach to reprogram human T cells with CRISPR-based genome targeting without the need for viral vectors^5^. However, many research and clinical applications still depend upon improved efficiency, cell viability, and generalizability of non-viral genome targeting across cell types^1–4^. We previously found that varying the relative concentrations of both Cas9 RNP and HDR template had significant effects on targeting efficiency and toxicity^5^. Here, we set out to optimize the interactions between the HDR template and stabilized RNPs to further improvement genome editing efficiency independent of cell type.

We devised a novel approach to promote nuclear entry of the template. Unlike previous efforts utilizing complex covalent linkages^7^, we attempted to recruit Cas9 RNPs with nuclear location sequences (NLS) to the HDR template by enhancing Watson-Crick interactions. CRISPR-Cas9 interacts specifically with both genomic and non-genomic dsDNA^8^, and nuclease-inactive dCas9 has been used in many applications to localize protein and RNA effectors to specific DNA sequences without cleaving the target sequence^9^. We therefore tested if we could enhance HDR by targeting a dCas9-NLS ‘shuttle’ to the ends of an HDR template by coding 20 bp Cas9 Target Sequences (CTS) at the ends of the homology arms. Indeed, CTS-modified HDR templates mixed with dCas9-NLS RNP did show mild improvements in HDR efficiency in primary human T cells (**Supplementary Fig. 1**) but required two distinct RNPs. These data encouraged us to search for a simplified method utilizing the same RNP to both cut a specified genomic site and recruit Cas9-NLS to HDR templates.

We hypothesized that a single catalytically-active Cas9-NLS RNP would suffice for both on-target genomic cutting and ‘shuttling’ if the HDR template was designed with truncated (16bp) Cas9 Target Sequences (tCTS) that enable Cas9 binding but not cutting^10^ (Fig. 1a and **Supplementary Fig. 2**). With the proper sequence orientation, the additional tCTS markedly improved knockin efficiency of a 1.5kb DNA sequence inserting a reprogrammed TCRα and TCRβ specificity at the endogenous *TRAC* (T-cell receptor α constant) locus (Fig. 1b and **Supplementary Fig. 2**). This tCTS shuttle system also improved genome targeting efficiencies across a variety of loci in different primary human T cell subsets (Fig. 1c,d and **Supplementary Fig. 3**). HDR templates with the tCTS achieved preferential targeting even in direct competition with unmodified dsDNA HDR templates simultaneously delivered to the same cells (Fig. 1e and **Supplementary Fig. 4**). Additionally, the tCTS shuttle improved efficiencies of bi-allelic and multiplexed targeting across different loci (Fig. 1f and **Supplementary Fig. 4**). The full HDR efficiency gains critically depended on the presence of an NLS in the Cas9 RNP, use of an on-target gRNA, and pre-incubation of the Cas9-NLS RNP with the tCTS-modified HDR template (Fig. 1g and **Supplementary Fig. 5**). Taken together these data demonstrate that coupling the HDR template with tCTS to a Cas9-NLS RNP enhances genome targeting efficiency without requiring modification of the protein or gRNA.

**Figure 1.**
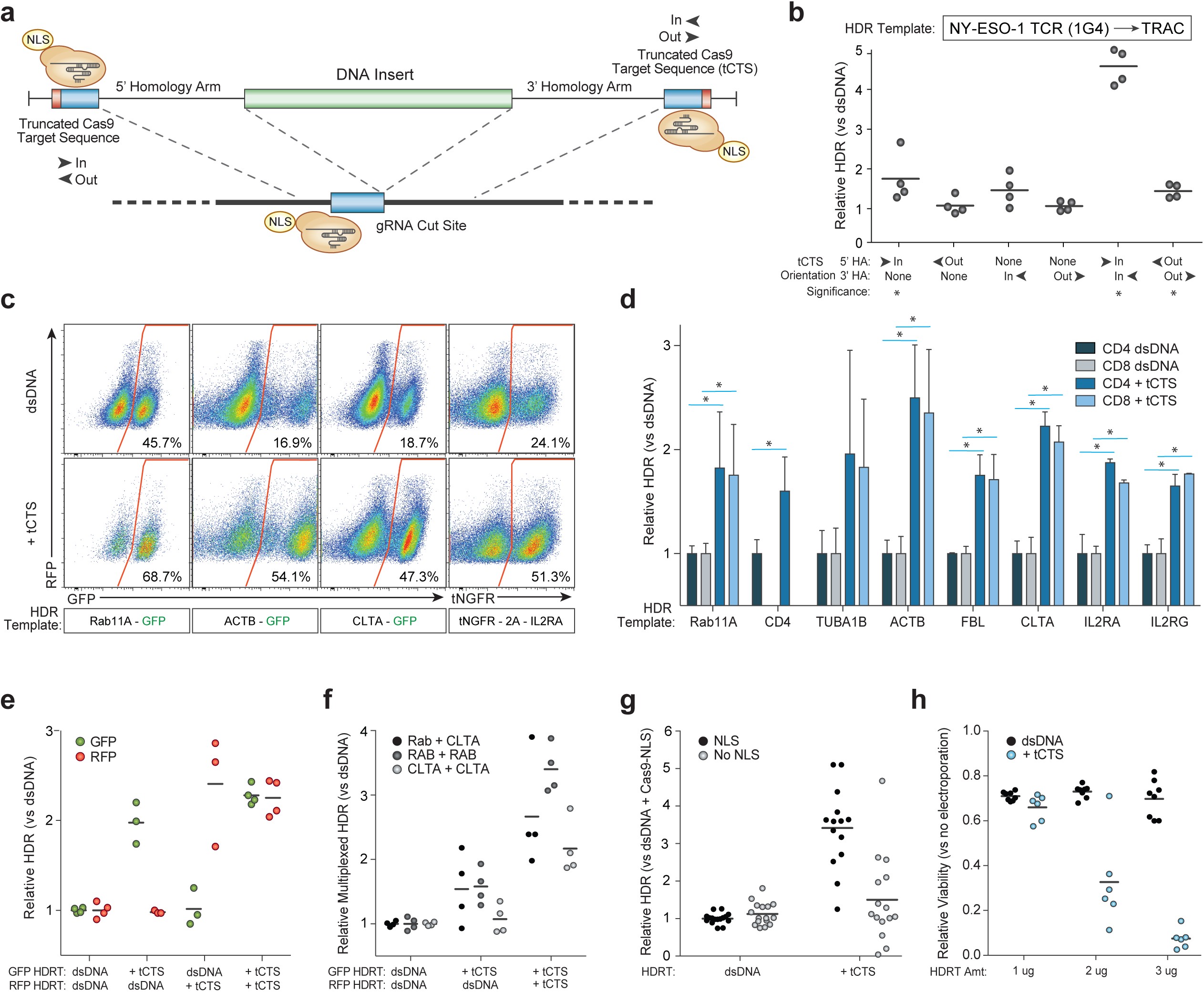
Truncated Cas9 Targets Sequences (tCTS) in HDR templates increase large non-viral knockin efficiency. (**a**) Enzymatically active Cas9-NLS RNPs can bind truncated Cas9 target sequences (tCTS) added to the ends of an HDR template. (**b**) “In” facing orientation of the tCTSs (PAM facing in towards the center of the inserted sequence vs “out” away from the insert) on the edges of both the 5’ and 3’ homology arms improved knockin efficiency of a new TCRα-TCRβ specificity at the endogenous *TRAC* locus. (**c**) Representative flow cytometry plots showed improved targeting efficiency across target genomic loci with the tCTS modifications compared to an unmodified dsDNA HDR template. (**d**) The tCTS modifications improved targeting efficiencies of large knockins across eight genomic loci tested in both CD4+ and CD8+ T cells. Note CD4-GFP expression was not observed at relevant levels in CD8+ T cells, as expected. (**e**) Multiplexed electroporation of GFP and RFP knockin templates to the *RAB11A* locus where neither, one, or both templates had a tCTS modification revealed direct competitive knockin advantage of ‘shuttle’ system compared to unmodified dsDNA template (technical replicates from n=2 donors). (**f**) The tCTS modification improved multiplexed dual knockin at different genomic loci as well as bi-allelic knockin at a single target locus (technical replicates from n=2 donors). (**g**) Full improvement of knockin efficiencies with the tCTS modifications were dependent on Cas9-NLS protein (multiple technical replicates from n=2 donors). (**h**) Decreased viability was seen with the tCTS modifications at lower DNA concentrations compared to unmodified dsDNA HDR template. The relative rates of HDR (**b, d, e, g**), multiplexed HDR (**f**), or viability (**h**) with the tCTS shuttle are displayed compared to unmodified dsDNA HDR template (**b, d-g**) or to no electroporation controls (**h**) in n=4 donors (**b, c, d**) or multiple technical replicates from n=2 donors (**e-h**); error bars indicate standard deviation. HDR efficiency was measured 4 days post electroporation and viability (total number of live cells relative to no electroporation control) at 2 days post electroporation.

Exogenous DNA (including HDR templates) can be cytotoxic at high concentrations^4,5,11^. We therefore assayed the effects of the RNP-HDR template interactions on cell viability. Gene targeting using tCTS-modified HDR templates improved efficiency, but we observed decreased cell viability at DNA doses lower than unmodified dsDNA HDR templates (Fig. 1h). Decreased viability was only observed with an on-target gRNA and pre-incubation of the RNP with HDR template, but did not entirely depend on the presence of a NLS on Cas9 (**Supplementary Fig. 5**) consistent with possible enhanced cytoplasmic delivery of DNA during electroporation due to the RNP-HDR template interaction.

In previous experiments, we observed that Cas9 RNP co-delivery could mitigate exogenous DNA toxicity (to unmodified plasmids or dsDNA template) in human T cells^5^. We therefore wondered whether optimizing RNP delivery could improve cell viability. We noted that the Cas9 protein itself appears to be only quasi-stable when complexed with guide RNA; a molar excess of protein (RNA to Cas9 protein molar ratio of <1.0) results in a milky opaque solution with rapid sedimentation (Fig. 2a) of poorly functional material (**Supplementary Fig. 6a**). Previous reports have suggested that RNP electroporation can be improved with addition of extra sgRNA or a non-homologous single strand DNA enhancer (ssODNenh)^12^. Excess guide RNA or addition of ssODNenh dispersed the Cas9 RNPs (**Supplementary Fig. 6b**) and boosted editing efficiency of electroporated RNPs (**Supplementary Fig. 6c**). However, combination of both guide RNA and ssODNenh in excess did not further improve editing (**Supplementary Fig. 6c**), suggesting a possible shared mechanism of action.

**Figure 2:**
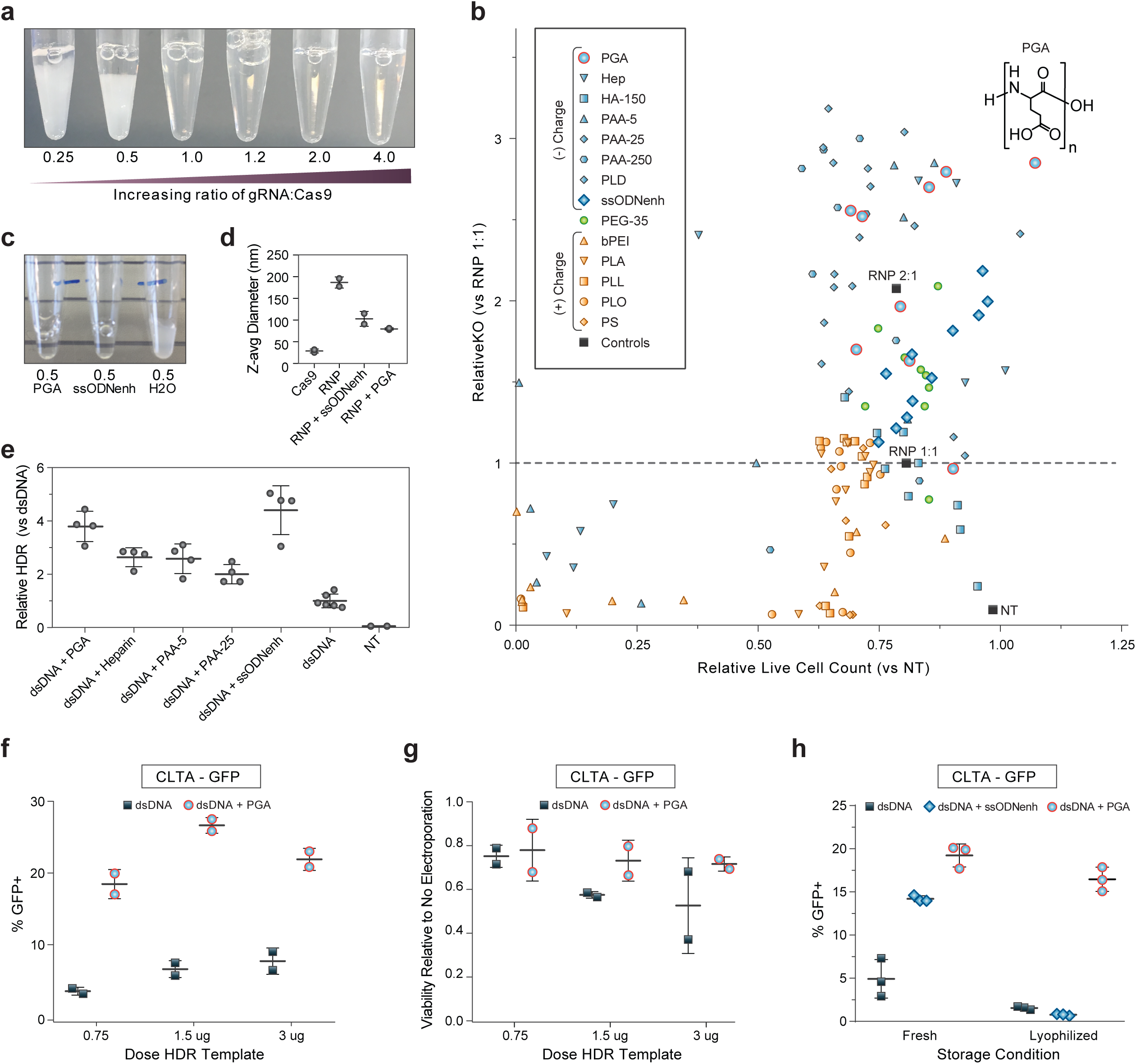
Stabilizing Cas9 RNP nanoparticles with anionic polymers improves editing outcomes. (**a**) Photograph 15 minutes after mixing gRNA and Cas9 protein incubated at 37C to form RNPs. Cas9 RNPs prepared at low molar ratio of gRNA:protein appeared cloudy and rapidly settled out of solution. (**b**) Multiple polymers were screened for the ability to enhance *CD4* gene knock-out editing when mixed with RNPs formulated at 1:1 gRNA:protein ratio then electroporated into primary human CD4+ T cells. Loss of surface CD4 expression at 3 days assessed by flow cytometry is normalized to unenhanced editing efficiency (RNP 1:1 without any additive) on the y-axis, and the live cell count is normalized to mock non-electroporated (NT) cells on the x-axis. Negatively charged polymers are shaded blue: poly(L-glutamic acid) (PGA), heparin sulfate (Hep), hyaluronic acid 150 kDa (HA-150), poly(acrylic acid) at 5 kDa (PAA-5) 25 kDa (PAA-25) or 250 kDa (PAA-250), poly(L-aspartic acid) (PLD), ssODNenh. Neutral polymers are shaded green: poly(ethylene glycol) 35 kDa (PEG-35), and positively charged polymers are shaded orange: polyethyleneimine 25 kDa (PEI), poly(L-arginine) 15-70 kDa (PLA), poly(L-lysine) 15-30 kDa (PLL), poly(L-ornithine) 30-70 kDa (PLO), and protamine sulfate (PS). PGA (blue circles outlined in red) with chemical structure shown inset above data point that corresponds to 100mg/mL concentration. Each polymer sample was tested at serial dilutions to avoid potential dose-dependent cytotoxicity falsely masking impact on editing efficiency, and each concentration is depicted as an individual point that is an average for two different blood donors. (**c**) Cas9 RNPs at 0.5 molar ratio of gRNA:protein (prepared same as in (a)) could be further dispersed with addition of PGA or ssODNenh, whereas dilution with water alone had no visible benefit. (**d**) PGA and ssODNenh stabilized and reduced the size of RNP nanoparticles. Cas9 RNPs prepared at 2:1 molar ratio of gRNA:protein alone (RNP) or mixed with PGA or ssODNenh were assessed for hydrodynamic particle size by dynamic light scattering. Z-average particle size is shown for n=2 independent preparations each averaged over ten repeated measurements (individual sample size distributions and peaks shown in Supplementary Fig. 8). (**e**) Multiple anionic polymers boosted knockin editing efficiency. Polymers mixed with Cas9 RNPs prepared at a 2:1 gRNA:protein ratio were further mixed with 1ug unmodified dsDNA HDR templates (targeting insertion of an N-terminal fusion of GFP to Rab11a), electroporated into CD4+ T cells, and editing efficiency were assessed by flow cytometry at day 3. The relative rates of HDR is displayed compared to unmodified dsDNA HDR template without enhancer. Two technical replicates shown for two different blood donors each. (**f**) PGA-stabilized Cas9 RNPs prepared at a 2:1 ratio gRNA:protein markedly improved knockin editing in primary human Bulk (CD3+) T cells targeting a C-terminal fusion of GFP to clathrin and (**g**) improve viability of electroporated cells (compared to untreated cells). Data shown as average from two different blood donors. (**h**) Cas9 RNPs prepared at a 2:1 ratio gRNA:protein without or with PGA or ssODNenh were mixed with 1ug of unmodified dsDNA HDR template targeting an N-terminal fusion of GFP to *RAB11A*, lyophilized overnight, stored dry at −80C, then later reconstituted in water prior to electroporating into primary human CD3+ (Pan) T cells. PGA-stabilized Cas9 nanoparticles were protected through lyophilization and reconstitution and retained activity for robust knockin editing. Three technical replicates shown for one blood donor. (**d-h**) Error bars indicate standard deviation.

We hypothesized that the polymeric and anionic nature of nucleic acids shields excess positively-charged residues of the Cas9 protein from nearby exposed portions of Cas9-bound gRNA, thus preventing aggregation and improving RNP particle stability. We therefore screened various commercially-available water-soluble biological and synthetic polymeric materials for the ability to also enhance electroporation-mediated Cas9 knock-out editing. Multiple different anionic polymers such as poly(glutamic acid) (PGA), poly(aspartic acid), heparin, and poly(acrylic acid) all enhanced editing efficiency in a dose-dependent manner without addition of ssODNenh or excess gRNA (Fig. 2b, **Supplementary Fig. 7**). The charge-neutral material poly(ethyleneglycol) (PEG) had only minimal impact on RNP activity; positively charged polymers poly(ethylenimine), protamine sulfate, poly-L-lysine, poly-L-ornithine, and poly-L-arginine all reduced editing efficiency (and viability) (Fig. 2b), thus establishing anionic charge as a key factor for RNP enhancement. Further, enhancement of RNP-based editing depended upon polymer chain length (**Supplementary Fig. 7**) similar to the reported length-dependence of ssODNenh^12^ and consistent with colloid-stabilizing biomaterials^13^. Comparison of particle sizes by dynamic light scattering revealed that Cas9 protein by itself is 10-15nm in diameter as expected for dispersed individual molecules, but aggregates of ~200+ nm size were formed when gRNA was added (Fig. 2d and **Supplemental Fig. 8**). However, the addition of either PGA or ssODNenh prevented aggregation into micron-sized particles and improved the size distribution of RNP nanoparticles to less than 100nm on average, (Fig. 2d) with peaks in the 20nm and 100-120nm ranges (**Supplementary Fig. 8**).

We thus tested if anionic polymers could also enhance non-viral knockin genome targeting. When mixed with Cas9 RNPs and a reduced concentration of an unmodified dsDNA template targeting integration of a Rab11a-GFP fusion, PGA was effective at enhancing knockin editing in primary T cells (Fig. 2e). PGA-stabilized RNP nanoparticles promoted efficiency gains in primary human T cells (Fig. 2f) and appeared to reduce the toxicity of higher doses of HDR template (Fig. 2g). These efficiency gains were independent of the order of PGA addition, guide RNA source, or Cas9 nuclease manufacturer (**Supplemental Figs. 9-12**). RNP stabilization with PGA (but not ssODNenh) also permitted lyophilization with retained knockout and knockin editing activity (Fig. 2h, **Supplementary Fig. 13**). PGA-stabilized RNP nanoparticles therefore enhance edited cell viability and could be stored dry enabling significant streamlining of gene modified cell manufacturing for research or clinical translation.

We next examined if combining the tCTS ‘shuttle’ and anionic polymers could further improve efficiency and viability of non-viral genome targeting. The tCTS-modified HDR templates with PGA-stabilized RNPs markedly enhanced efficiency and viability in primary human T cells across multiple genomic loci (Fig. 3a). Importantly, the dose-dependent toxicity observed with the tCTS-modified HDR template (Fig. 1h) was mitigated by PGA-stabilized RNPs and improved the recovery of viable T cells edited at a variety of endogenous loci (Fig. 3b). Increased efficiency and cell recovery were consistent across human blood donors and were most profound at lower concentrations of HDR template DNA (**Supplementary Fig. 14**). The combined tCTS-modified HDR template and PGA-stabilized nanoparticle system also retained activity through freeze-thaw cycles and the lyophilization process (**Supplementary Fig. 15**).

**Figure 3:**
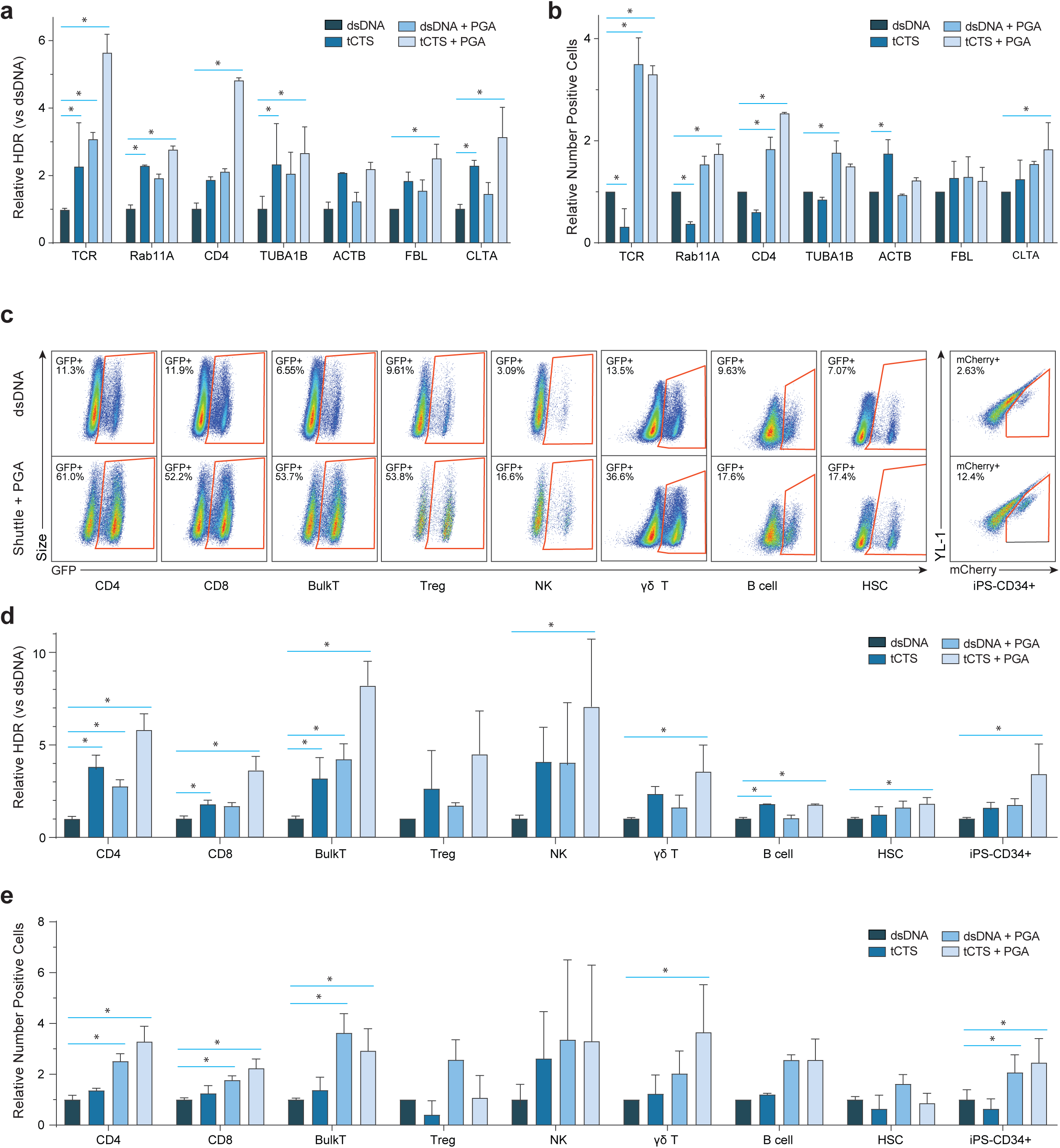
PGA-stabilized Cas9 RNP and tCTS-modified-HDR templates improved knockin gene editing outcomes across a variety of genetic loci and clinically-relevant immune cell types. (**a-b**) Cas9 RNPs were prepared at a 2:1 ratio gRNA:protein with or without PGA polymer and mixed with high doses (2-4 ug) of regular dsDNA or tCTS-modified HDR template targeting knockin at multiple genomic loci: transgenic NY-ESO 1 tumor antigen T cell receptor into the *TRAC* locus, or GFP fusion at the N- or C- of *RAB11A, CD4, TUBA1B, ACTB, FBL, or CLTA* genes. The combination of PGA-stabilized Cas9 RNP nanoparticles and ‘shuttle’ tCTS-modified-HDR template both improved relative frequency of HDR (**a**) and resulted in higher yield of successfully edited cells (**b**). (**c-e**) Cas9 RNPs were prepared at 2:1 ratio gRNA:protein with or without PGA polymer and mixed with low doses (0.5 – 1 ug) of unmodified dsDNA or ‘shuttle’ tCTS-modified HDR templates targeting knockin of GFP or mCherry to the N-terminus of Clathrin. The PGA-stabilized Cas9 RNP nanoparticles and tCTS-modified HDR templates improved editing efficiency in a variety of primary human immune cell types as visualized in representative flow cytometry plots (after gating for live cells and respective cell type-specific surface markers) (**c**) or expressed as relative frequency of GFP or mCherry+ positive cells (**d**), and resulted in higher yield of number of successfully edited cells (**e**). The relative rates of HDR (**a,d**) or edited cell recovery (**b,e**) is displayed compared to unmodified dsDNA HDR template without enhancer. Error bars indicate standard deviation; (*) indicates significant difference compared to control unmodified dsDNA HDR template without enhancer with p<0.05 (ANOVA with Bonferroni’s multiple comparisons test correction) for each given gene locus (**a-b**) or cell type (**d-e**).

Encouraged by these results in T cells, we applied the combined system to enable non-viral genome targeting in a wider set of therapeutically relevant primary human hematopoietic cells. Using a Clathrin-GFP fusion template, which should be expressed in all tested cell types, we consistently observed improved editing efficiencies with the combined system (Fig. 3c-e). Bulk CD3+ T cells, purified CD4+ T cells or CD8+ T cells, and purified CD127low CD25+ CD4+ regulatory T cells (Tregs) all achieved a similarly high knockin efficiency of up to > 50% of cells, with 3-8x increase in the percentage of knockin edited cells at reduced HDR template concentrations (Fig. 3c,d). While standard RNP and unmodified HDR template achieved only minimal knockin in isolated primary human NK cells or B cells, the combined system resulted in over 15% transgene-positive cells and a 2-5-fold increase in edited cell yield (Fig. 3c-e). In γδ-T cells the combined system exhibited improved editing efficiency from ~5% to ~28%, and 5-6 fold improved edited cell recovery compared to a standard RNP and unmodified HDR template. Finally, we were able to express large transgene insertions in over 15% of human HSPCs without a viral vector, in both primary mobilized peripheral blood- and iPS-derived^6^ CD34+ HSPCs, with a marked viability boost and 2-3x increased yield of edited cells. The combined non-viral system was thus demonstrated to be effective in diverse human hematopoietic cell types.

Together, PGA as a RNP nanoparticle-stabilizing enhancer and the tCTS-modified HDR template enabled high percentage editing with improved edited cell yields in a variety of primary hematopoietic cell types opening the door to next-generation adoptive cell therapies beyond T cells. The combined nanoparticle-tCTS template system is a novel platform to explore gene function in clinically-relevant cell types that had been previously challenging to genome modify. The formation of PGA-stabilized RNP nanoparticles and the utilization of PCR primers to introduce tCTS modifications to HDR templates are both methods that can be rapidly adapted to any existing Cas9 RNP-based editing protocol. Notably, marked improvements in large gene targeting to endogenous loci were achieved without further optimizing cell cycle dynamics, small molecule modulation of DNA repair machinery, or specialized chemistries; these complementary strategies may eventually offer additional efficiency gains. Further optimization of polymers or tCTS ‘shuttle’ configurations could offer even further improvements. Our data demonstrate a technically simple system that greatly enhances the capabilities of Cas9 RNP-mediated non-viral genome targeting in primary human hematopoietic cells and has direct translational potential for research, biotechnology, and clinical endeavors.

## ACKNOWLEDGEMENTS

We thank members of the Marson lab, Chris Jeans and the QB3 MacroLab, and Aaron Dolor for suggestions and technical assistance. This research was supported by NIH grants DP3DK111914-01, R01DK1199979 and P50GM082250 (A.M.), a grant from the Keck Foundation (A.M.), gifts from Jake Aronov and Barbara Bakar (A.M.), a gift from the UCSF Diabetes Center, a gift from the Jeffrey Modell Foundation, and a National Multiple Sclerosis Society grant (A.M.; CA 1074-A-21). D.N.N was supported by the UCSF Biology of Infectious Diseases Training Program (T32A1007641) and the CIS CSL Behring Fellowship Award. T.L.R. was supported by the UCSF Medical Scientist Training Program (T32GM007618), the UCSF Endocrinology Training Grant (T32 DK007418), and the NIDDK (F30DK120213). A.M. holds a Career Award for Medical Scientists from the Burroughs Wellcome Fund, is an investigator at the Chan Zuckerberg Biohub and has received funding from the Innovative Genomics Institute (IGI) and the Parker Institute for Cancer Immunotherapy (PICI). The UCSF Flow Cytometry Core was supported by the Diabetes Research Center grant NIH P30 DK063720.

## AUTHOR CONTRIBUTIONS

D.N.N., T.L.R., and A.M. designed the study. T.L.R. conceived of the template ‘shuttle’ system and performed all ‘shuttle’ optimization. D.G. suggested the use of truncated Cas9 Target Sequences. D.N.N. conceived of the polymer stabilization of RNPs system and performed all polymer optimizations. D.N.N., T.L.R., J.L., P.A.C., M.R.M., L.T.V., V.T., R.A., D.G., E.S., J.A.B., J.M.P., and F.C.S. contributed to the design and completion of experiments combining the shuttle and polymer systems in additional primary cell types. D.N.N., T.L.R., and A.M. wrote the manuscript with input from all authors.

## METHODS

### Cell Culture

Primary adult cells were obtained from healthy human donors from leukoreduction chamber residuals after Trima Apheresis (Vitalant, formerly Blood Centers of the Pacific) or from freshly drawn whole blood under a protocol approved by the UCSF IRB (BU101283). Peripheral blood mononuclear cells (PBMCs) were isolated by Ficoll-Paque (GE Healthcare) centrifugation using SepMate tubes (STEMCELL, per manufacturer’s instructions). Specific lymphocytes were then further isolated by magnetic negative selection using an EasySep Human B Cell, CD4+ T Cell, Bulk (CD3+) T Cell, CD8+ T cell, CD4+ CD127low CD25+ Regulatory T Cell, Gamma/Delta T Cell, or NK Cell Isolation kit (STEMCELL, per manufacturer’s instructions).

Isolated CD4+, CD8+, CD3+ (Bulk T Cells), Regulatory (CD25hiCD127low), or γδ-T cells were activated and cultured for 2 days at 0.5 to 1.0 million cells/mL in XVivo15 medium (Lonza) with 5% Fetal Bovine Serum, 50 mM 2-mercaptoethanol, 10 mM N-Acetyl L-Cystine, anti-human CD3/CD28 magnetic Dynabeads (ThermoFisher) at a beads to cells concentration of 1:1, and a cytokine cocktail of IL-2 at 300 U/mL (UCSF Pharmacy), IL-7 at 5 ng/mL (ThermoFisher), and IL-15 at 5 ng/mL (ThermoFischer). Activated T cells were harvested from their culture vessels and Dynabeads were removed by placing cells on an EasySep cell separation magnet (STEMCELL) for 5 minutes. Isolated B cells were cultured in IMDM (ThermoFischer) with 5% Fetal Bovine Serum, 100 ng/mL MEGACD40L (Enzo), 1000 ng/mL CpG (InvivoGen), 500 U/mL IL-2 (UCSF Pharmacy), 50 ng/mL IL-10 (ThermoFischer), and 10 ng/mL IL-15 (ThermoFischer). Freshly isolated NK cells were cultured in Xvivo15 medium (Lonza) with 5% Fetal Bovine Serum, 50 mM 2-mercaptoethanol, 10 mM N-Acetyl L-Cystine, together with IL-2 (at 1000 U/mL) and MACSiBead Particles pre-loaded with anti-human CD335 (NKp46) and anti-human CD2 antibodies (Miltenyi Biotech). Primary adult peripheral blood G-CSF-mobilized CD34+ hematopoietic stem cells were purchased from StemExpress Inc and cultured at 0.5e6 cells/mL in SFEM II media supplemented with CC110 cytokine cocktail (STEMCELL) for two days prior to electroporation. Induced pluripotent stem-cells were generated and differentiated into CD34+ HSPCs as previously described^6^ then cultured in SFEM media (STEMCELL) supplemented with IL-2 at 10ng/mL, IL-6 at 50ng/mL, SCF at 50ng/mL, FLT-3L at 50ng/mL, and TPO at 50ng/mL (Peprotech).

### RNP Formulation with Polymers

Cas9 RNPs were formulated immediately prior to electroporation, except when frozen or lyophilized as indicated. Synthetic crRNA (with guide sequence listed in Supplementary Table 1) and tracrRNA were chemically synthesized (Edit-R, Dharmacon Horizon or Integrated DNA Technologies (IDT)),resuspended in 10 mM Tris-HCL (7.4 pH) with 150 mM KCl or IDT duplex buffer at a concentration of 160 µM, and stored in aliquots at −80C. To make gRNA, aliquots of crRNA and tracrRNA were thawed, mixed 1:1 by volume, and annealed by incubation at 37C for 30 min to form an 80 µM gRNA solution. For comparison, chemically modified gRNA was purchased from Synthego and resuspended according to manufacturer’s protocol. Cas9-NLS, dCas9-NLS, or Cas9 without NLS was purchased from the UC Berkeley QB3 MAcrolab; HiFiCas9 was purchased from IDT. To make RNPs, gRNA was then further diluted in buffer first, or directly mixed 1:1 by volume with 40 µM Cas9-NLS protein to achieve the desired molar ratio of gRNA:Cas9 (2:1 ratio unless otherwise stated). Unless otherwise stated, final dose of RNP per nucleofection was 50pmol on a Cas9 protein basis.

For initial screening, polymers were purchased dry (see Supplementary Table 2) and resuspended to 100mg/mL in water except as noted, passed through a 0.2 um syringe filter, and stored at −80C prior to use. The ssODNenh electroporation enhancer (with sequence listed in Supplementary Table 2) was synthesized (IDT) and resuspended to 100 uM in water. Serial dilutions of polymers or ssODNenh were made in water, then mixed 1:1 volume with preformed RNPs. For subsequent knockin experiments, 15-50 kDa poly(L-glutamic acid) (Sigma) was resuspended to 100mg/mL in water, sterile filtered, and mixed with freshly-prepared gRNA at 0.8:1 volume ratio prior to complexing with Cas9 protein for final volume ratio gRNA:PGA:Cas9 of 2:1.6:1.

Long double-strand HDR templates encoding various gene insertions (see Supplementary Table 1) and 300-350bp homology arms were synthesized as GeneBlocks (IDT) and cloned into a pUC19 plasmid, which then served as a template for generating a PCR amplicon. Specific PCR primers targeting the left and right homology arms and with additional described Cas9 Target Sequences (CTS) (see Supplementary Figure 2) were synthesized (IDT) without chemical modifications. Amplicons were generated as previously described^5^ with KapaHiFi polymerase (Kapa Biosystems), purified by SPRI bead cleanup, and resuspended in water to 0.5 to 2 ug/uL measured by light absorbance on a NanoDrop spectrophotometer (ThermoFisher). HDR templates were mixed and incubated with RNPs for at least 5 minutes prior to mixing with and electroporating into cells.

RNP particle size was measured by dynamic light scattering dispersed in PBS on a Zetasizer Nano ZS (Malvern Panalytical). For RNP lyophilization, freshly prepared RNPs premixed with PGA or ssODNenh and HDR templates were diluted 1:1 v:v in 50mM trehalose, flash frozen in a liquid nitrogen bath then immediately dried on a Labconco Freeze Dry System-Freezone 4.5 lyophilizer for 24 hours, and stored at −80C until use. Dry RNP was resuspended in water to achieve the original concentration, incubated for 5 minutes at 37C, the mixed with cells for electroporation.

Immediately prior to electroporation in a Lonza 4D 96-well format Nucleofector (Lonza), cells were centrifuged for 10 minutes at 90g, media aspirated, and resuspended in the electroporation buffer P3 (Lonza) using 17-20 µL buffer per 0.5 to 1.0 e6 cells. T cells, NK cells, and B cells were electroporated with pulse code EH-115; primary HSPCs with pulse code ER-100, and iPS-derived CD34 HSPCs with pulse code EY-100. Immediately after electroporation, cells were rescued with addition of 80uL of growth media directly into the electroporation well, incubated for 10-20 minutes, then removed and diluted to 0.5-1e6 cells/mL in growth media. Additional fresh growth media and cytokines were added every 48 hours.

At 3-5 days post electroporation (except for NK cells collected at 5-7 days), cells were collected for staining and flow cytometry analysis. Briefly, cells were stained for cell type-specific surface markers and live-dead discrimination (see list of antibodies in Supplementary Table 3) then analyzed on an Attune NxT Flow Cytometer with automated 96-well sampler (ThermoFisher) sampling a defined volume (between 50-150uL per well) to obtain quantitative cell counts. Cytometry data was processed and analyzed using FlowJo software (BD Biosciences).

