## Supplemental Figures and Tables 2-3 for "A Cas9 nanoparticle system with truncated Cas9 target sequences on DNA repair templates enhances genome targeting in diverse human immune cell types"

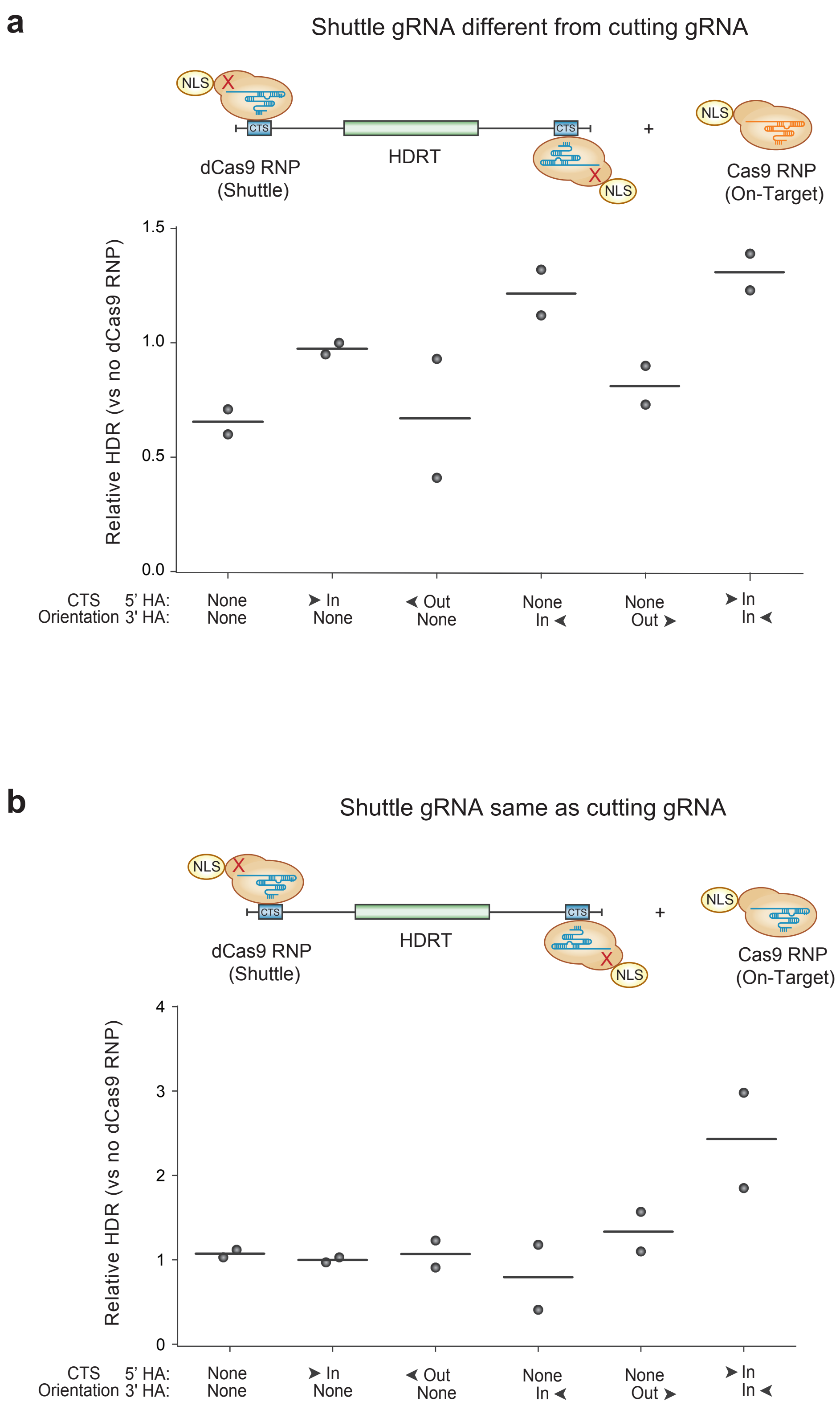
a
Shuttle gRNA different from cutting gRNA
NLS
CTS
HDRT
dCas9 RNP
(Shuttle)
Cas9 RNP
(On-Target)
Relative HDR (vs no dCas9 RNP)
1.5
1.0
0.5
0.0
CTS Orientation 5' HA:
3' HA:
None None
In None
Out None
None In
None Out
In In
b
Shuttle gRNA same as cutting gRNA
NLS
CTS
HDRT
dCas9 RNP
(Shuttle)
Cas9 RNP
(On-Target)
Relative HDR (vs no dCas9 RNP)
4
3
2
1
0
CTS Orientation 5' HA:
3' HA:
None None
In None
Out None
None In
None Out
In In

**Supplementary Fig. 1: dCas9 complexing to DNA HDR templates moderately improved knockin efficiencies.**

(**a**) Initial testing of a dCas9 shuttle. A full 20bp gRNA target sequence was added to the edge of a dsDNA HDR template that inserts a new TCR specificity (1G4 TCR clone, specific for the NY-ESO1 cancer antigen) at the endogenous TCR locus[5]. A dCas9 RNP was formed with a gRNA specific for the sequence attached to the ends of the HDR template, incubated with the dsDNA HDR template for 5 min at room temperature, and then electroporated into primary human T cells along with a separate Cas9 RNP with a gRNA specific for the target locus, in this case the TCR alpha constant region exon 1. The orientation of the gRNA target sequence added to the ends of the HDR template appeared to matter, with an “In” facing orientation (PAM sequence towards the center of the insert) on both the 5’ and 3’ homology arms showed the greatest improvement in knockin efficiency.

(**b**) The dCas9 shuttle RNP could be complexed with the same gRNA sequence as the on-target cutting Cas9 RNP by appending the same gRNA target sequence as in the on-target genomic locus to the ends of the HDR template. Similarly, an “In” facing orientation of the sequence on both edges of the homology arm were important for improved knockin efficiency.

The relative rates of HDR are displayed at four days post electroporation when including the dCas9 ‘shuttle’ RNP compared to electroporation of the unmodified HDR template and Cas9 for n=2 donors (**a, b**).

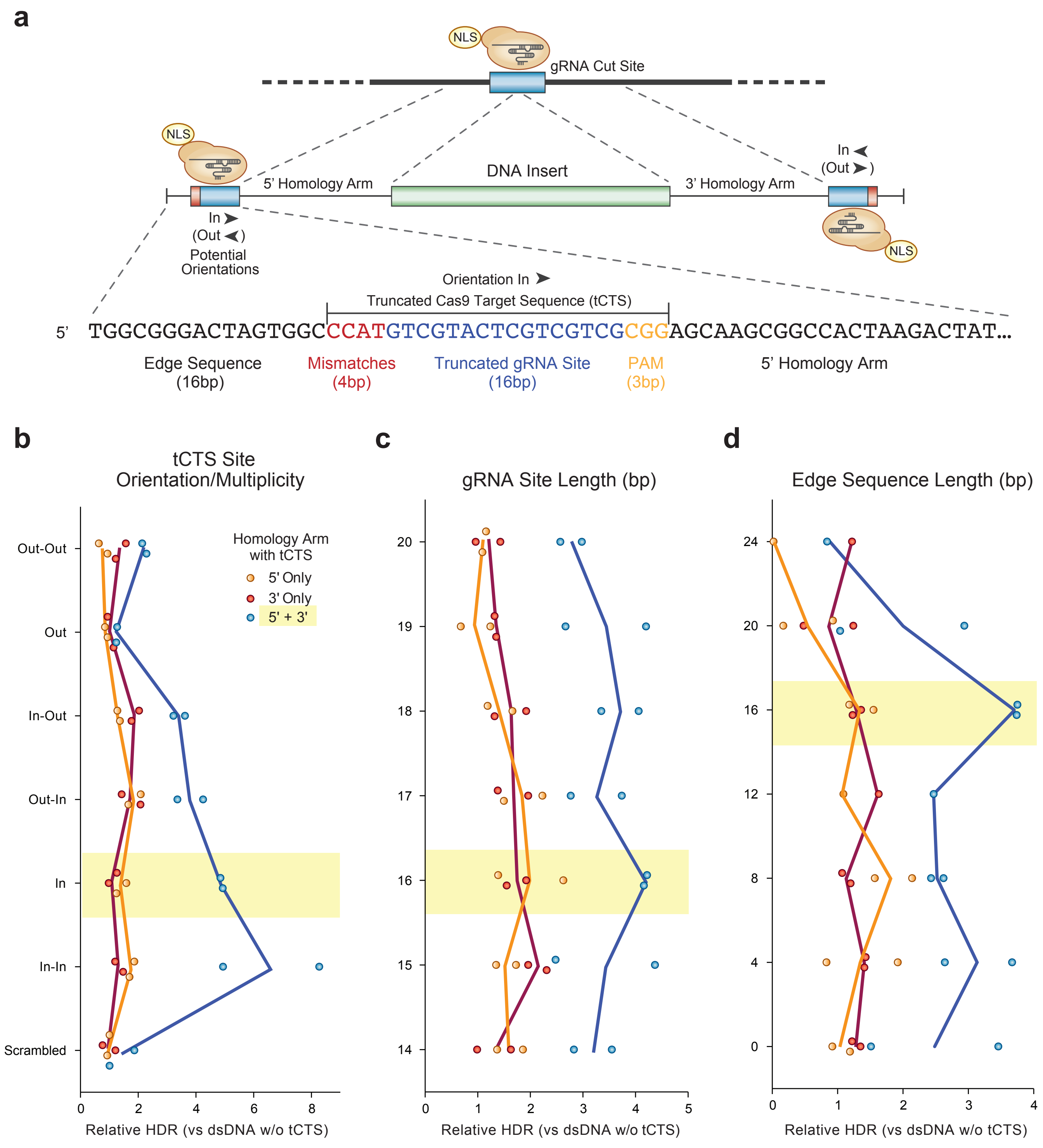
a
NLS
gRNA Cut Site
NLS
5' Homology Arm
DNA Insert
3' Homology Arm
In ➤
(Out ➤)
In ➤
(Out ➤)
Potential Orientations
NLS
Orientation In ➤
Truncated Cas9 Target Sequence (tCTS)
5' TGGCGGGACTAGTGGCCCATGTCGTACTCGTCGTCGCGGAGCAAGCGGCCACTAAGACTAT...
Edge Sequence (16bp)
Mismatches (4bp)
Truncated gRNA Site (16bp)
PAM (3bp)
5' Homology Arm
b
tCTS Site Orientation/Multiplicity
Out-Out
Out
In-Out
Out-In
In
In-In
Scrambled
Homology Arm with tCTS
5' Only
3' Only
5' + 3'
0 2 4 6 8
Relative HDR (vs dsDNA w/o tCTS)
c
gRNA Site Length (bp)
20 19 18 17 16 15 14
0 1 2 3 4 5
Relative HDR (vs dsDNA w/o tCTS)
d
Edge Sequence Length (bp)
24 20 16 12 8 4 0
0 1 2 3 4
Relative HDR (vs dsDNA w/o tCTS)

**Supplementary Fig. 2: Optimization of Cas9 'shuttle' end modification DNA sequence parameters.**

(**a**) DNA sequence design for the Cas9 'shuttle' system. A short DNA sequence can be easily added to the edges of a dsDNA HDR template by PCR. To mediate binding of a Cas9-NLS RNP to the HDR template but prevent cutting, a truncated 16 bp Cas9 target sequence is added to either or both of the 5' and 3' homology arms, along with a 3 bp PAM sequence (NGG for spCas9) and 4 bp of gRNA mismatches. An additional 16 bps of random DNA edge spacer sequence is added outside of the gRNA binding site. The gRNA site can have two orientations, one "In" facing with the PAM directed towards the center of the HDR template, and another "Out" facing with the PAM directed towards the edge of the HDR template.

(**b**) Optimization of the orientation and multiplicity of the Cas9 'shuttle' DNA sequence. Single or multiple 16 bp gRNA binding sites with PAMs were added to the 5' homology arm, 3' homology arm, or both homology arms of an HDR template that replaces the endogenous T cell receptor with a new specificity (insertion size ~1.5kb). Inwards facing orientations on both edges of the HDR template showed the greatest improvements in HDR efficiency, either with a single copy of the tCTS site ("In") or two copies in parallel ("In-In"). For simplicity a single copy was used in all subsequent experiments.

(**c**) Optimization of the length of the truncated gRNA binding site. Note that a full-length gRNA was used in all experiments, while the length of the CTS binding site varied. A DNA sequence with 16 base pairs matching the gRNA sequence proved optimal, corresponding to previously demonstrated gRNA target site lengths that mediate Cas9 binding but not cutting[8]. Again, presence of the tCTS-binding site on both homology arms increased HDR efficiency.

(**d**) Optimization of the edge spacer length showed that a 16 bp random DNA sequence outside of the tCTS yielded the greatest improvements in HDR efficiency. Maximal gains in HDR were again dependent on presence of the Cas9 'shuttle' sequence on both homology arms.

The relative rates of HDR (1.5 kb knockin) at the TCR locus with the indicated 'shuttle' sequence modifications compared to unmodified dsDNA HDR template for n=2 donors are displayed (**a, b**).

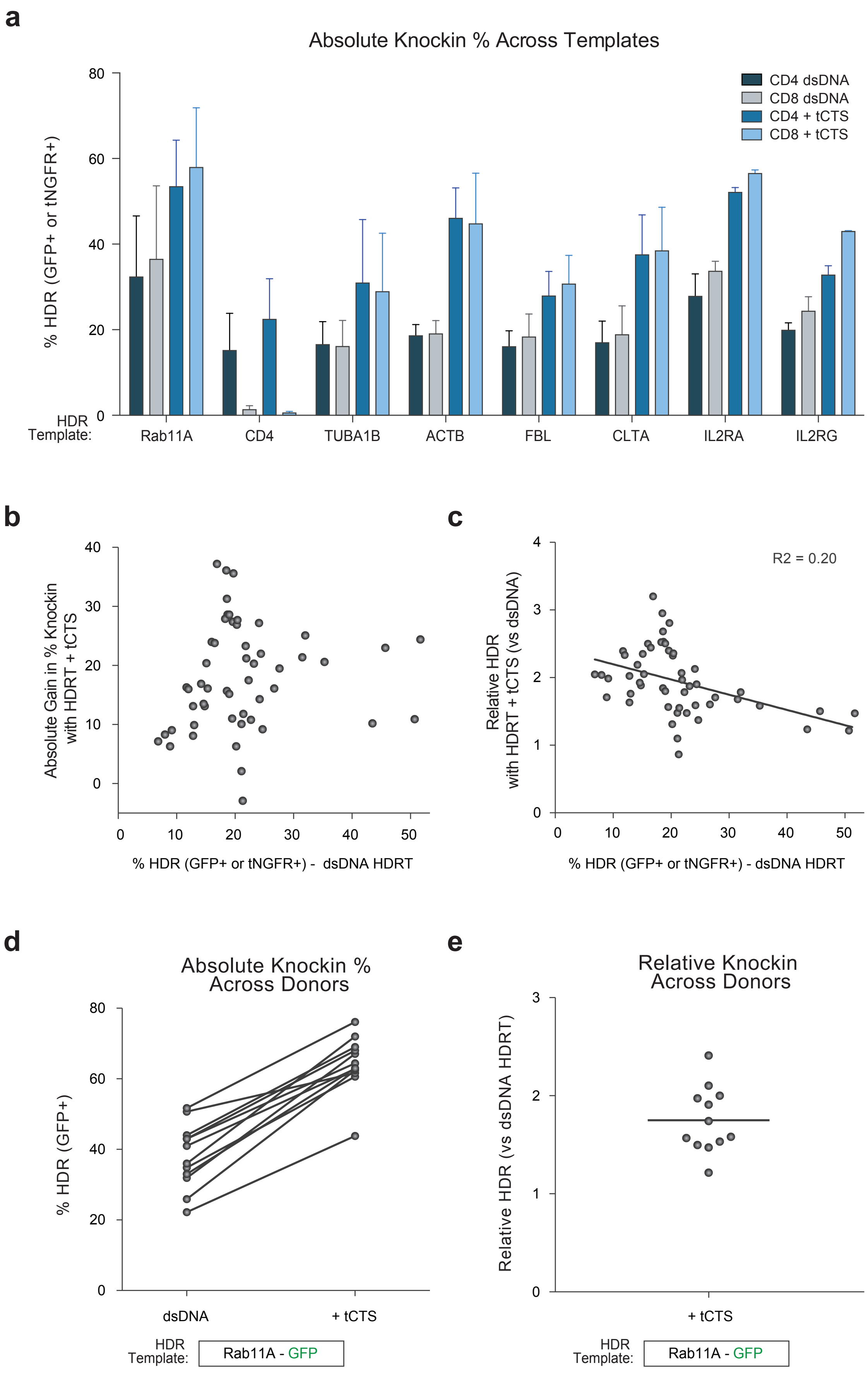
a
Absolute Knockin % Across Templates
% HDR (GFP+ or tNGFR+)
80
60
40
20
0
CD4 dsDNA
CD8 dsDNA
CD4 + tCTS
CD8 + tCTS
HDR Template:
Rab11A
CD4
TUBA1B
ACTB
FBL
CLTA
IL2RA
IL2RG
b
Absolute Gain in % Knockin with HDRT + tCTS
40
30
20
10
0
0
10
20
30
40
50
% HDR (GFP+ or tNGFR+) - dsDNA HDRT
c
Relative HDR with HDRT + tCTS (vs dsDNA)
R2 = 0.20
4
3
2
1
0
0
10
20
30
40
50
% HDR (GFP+ or tNGFR+) - dsDNA HDRT
d
Absolute Knockin % Across Donors
% HDR (GFP+)
80
60
40
20
0
dsDNA
+ tCTS
HDR Template:
Rab11A - GFP
e
Relative Knockin Across Donors
Relative HDR (vs dsDNA HDRT)
3
2
1
0
+ tCTS
HDR Template:
Rab11A - GFP

**Supplementary Fig. 3: Cas9 'shuttle' tCTS HDR template modifications improved large knockin efficiency across donors and genomic target loci.**

(**a**) Percentage of primary human CD4 or CD8 T cells expressing a knocked in GFP or tNGFR after non-viral genome targeting of eight different genomic sites with and without Cas9 'shuttle' tCTS-modifications to the dsDNA HDR template. Across all eight tested loci the Cas9 shuttle improved knockin percentages in both cell types. Note that knockin of a GFP to the CD4 locus did not show GFP expression in CD8 T cells.

(**b**) Aggregation of data across donors and target sites shows that the tCTS modification enabled large increases in the % of cells with an observed large knockin of GFP or tNGFR across initial knockin efficiencies with an unmodified dsDNA HDR template.

(**c**) Relative improvement in HDR efficiency is dependent on the initial knockin efficiency with an unmodified dsDNA HDR template. At lower initial knockin efficiencies, the relative gain in HDR with the Cas9 shuttle was higher than at target sites where the unmodified template efficiency was higher. Linear regression with a goodness of fit $R^2$ value of 0.20 is overlaid.

(**d**) In a cohort of 12 healthy primary T cell donors, all donors showed an improved knockin efficiency with the Cas9 'shuttle' compared to unmodified dsDNA HDR template for knockin of a GFP fusion at the *RAB11A* locus. Lines connect individual donor values.

(**e**) Relative improvement in HDR efficiency at the *RAB11A* locus with a Cas9 shuttle across 12 donors.

Knockins were assayed 4 days after electroporation in n=4 (**a-c**) or n=12 (**d-e**) blood donors.

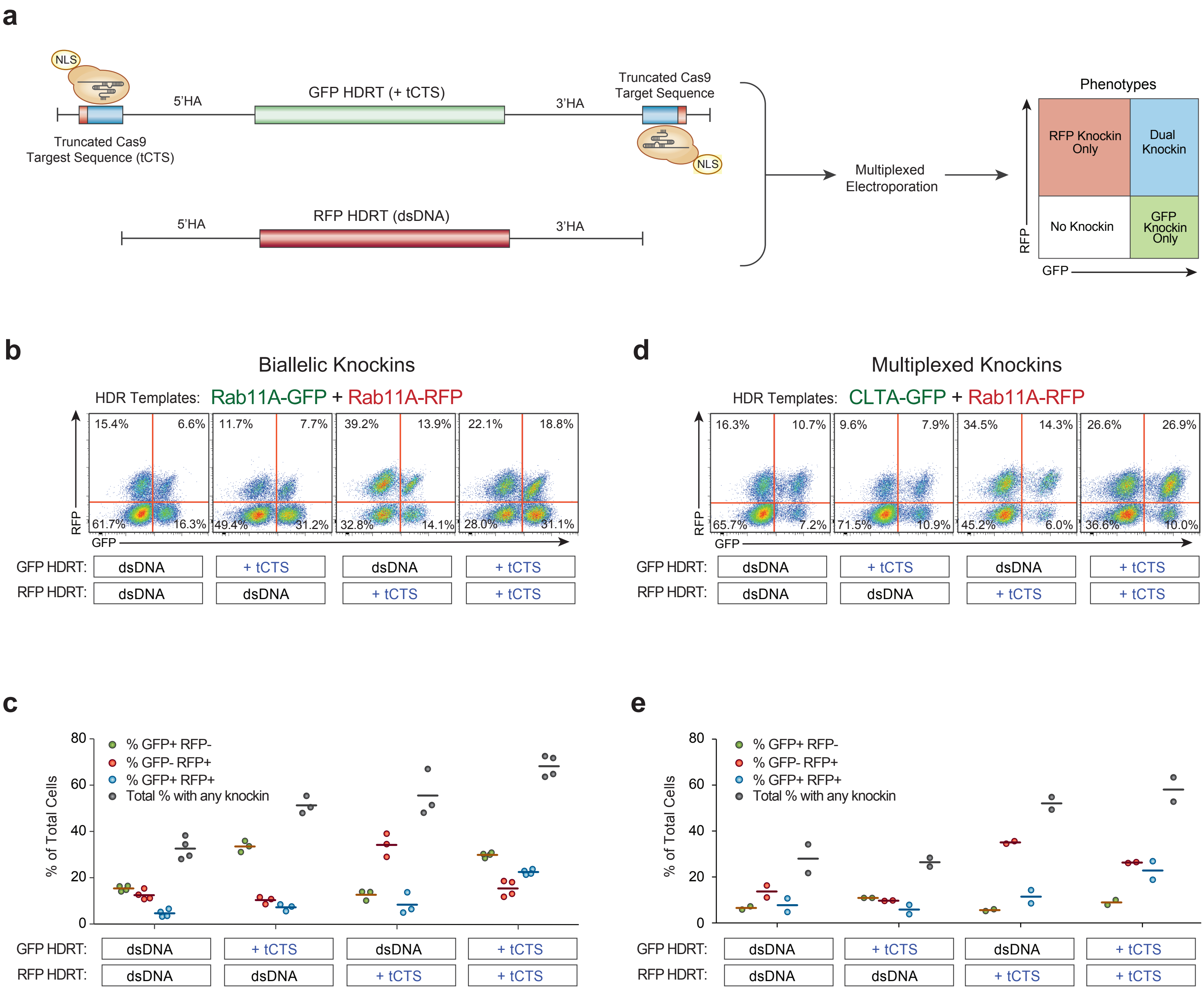
a
NLS
5'HA
GFP HDRT (+ tCTS)
3'HA
Truncated Cas9
Target Sequence
Truncated Cas9
Targest Sequence (tCTS)
NLS
5'HA
RFP HDRT (dsDNA)
3'HA
Multiplexed
Electroporation
Phenotypes
RFP Knockin Only
Dual Knockin
No Knockin
GFP Knockin Only
RFP
GFP
b
Biallelic Knockins
HDR Templates: Rab11A-GFP + Rab11A-RFP
15.4% 6.6% 61.7% 16.3%
11.7% 7.7% 49.4% 31.2%
39.2% 13.9% 32.8% 14.1%
22.1% 18.8% 28.0% 31.1%
RFP
GFP
GFP HDRT: dsDNA + tCTS dsDNA + tCTS
RFP HDRT: dsDNA dsDNA + tCTS + tCTS
d
Multiplexed Knockins
HDR Templates: CLTA-GFP + Rab11A-RFP
16.3% 10.7% 65.7% 7.2%
9.6% 7.9% 71.5% 10.9%
34.5% 14.3% 45.2% 6.0%
26.6% 26.9% 36.6% 10.0%
RFP
GFP
GFP HDRT: dsDNA + tCTS dsDNA + tCTS
RFP HDRT: dsDNA dsDNA + tCTS + tCTS
c
% of Total Cells
80
60
40
20
0
% GFP+ RFP-
% GFP- RFP+
% GFP+ RFP+
Total % with any knockin
GFP HDRT: dsDNA + tCTS dsDNA + tCTS
RFP HDRT: dsDNA dsDNA + tCTS + tCTS
e
% of Total Cells
80
60
40
20
0
% GFP+ RFP-
% GFP- RFP+
% GFP+ RFP+
Total % with any knockin
GFP HDRT: dsDNA + tCTS dsDNA + tCTS
RFP HDRT: dsDNA dsDNA + tCTS + tCTS

**Supplementary Fig. 4: Improved bi-allelic and multiplexed knockin efficiency with Cas9 shuttle.**

(**a**) Diagram of multiplexing experiments with two HDR templates, encoding either a GFP or RFP integration, simultaneously electroporated into the same cells. Neither, one, or both HDR templates had a Cas9 shuttle DNA sequence added to the ends of the homology arms, and the percentages of single and dual knockin positive cells were assayed by flow cytometry.

(**b**) Representative flow cytometric plots for bi-allelic knockin of GFP and RFP to the *RAB11A* housekeeping gene. When one HDR template possessed tCTS 'shuttle' modifications, the knockin rates for that fluorescent protein (either GFP or RFP) increased, whereas the knockin rates for the other fluorescent protein remained largely unaffected. Knockins with both templates possessing Cas9 shuttle sequences increased the percentage of bi-allelic targeting. Note that the percentage of GFP+RFP+ cells is only approximately half of the true bi-allelic knockin percentage, as knockins of either GFP or RFP at both alleles still yields a single positive at the protein level[5].

(**c**) Quantification of percent knockin efficiency across bi-allelic knockin experiments.

(**d**) Representative flow cytometric plots for multiplexed knockin experiments at two genomic loci, the *CLTA* locus encoding Clathrin and the *RAB11A* housekeeping gene. Again, the greatest improvements in multiplexed knockin efficiency were observed when both templates possess the Cas9 shuttle sequence.

(**e**) Quantification of percent knockin efficiency across multiplexed knockin experiments.

Knockins were assayed 4 days after electroporation in n=2 (**b-e**) blood donors.

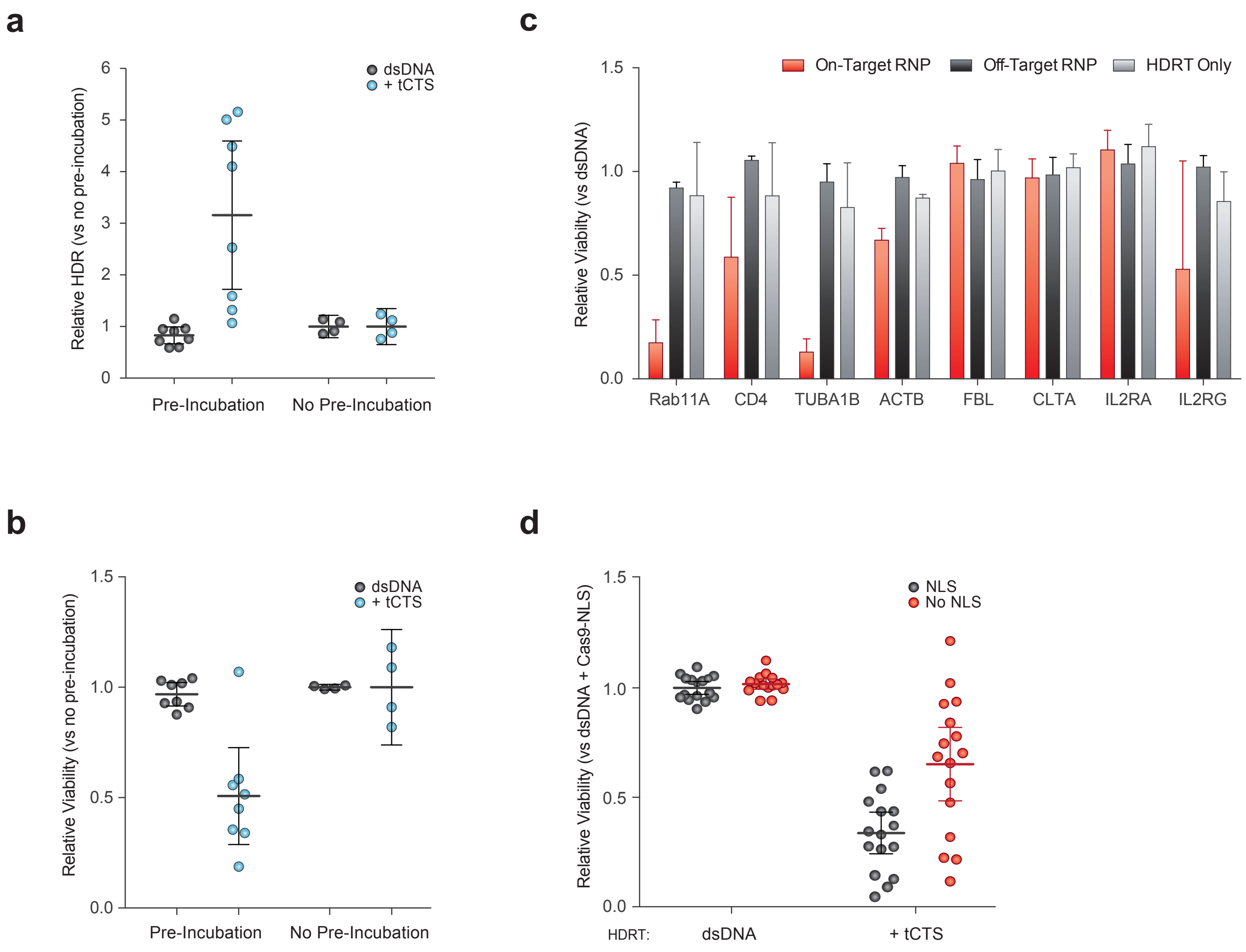
a
dsDNA
+ tCTS
Relative HDR (vs no pre-incubation)
Pre-Incubation
No Pre-Incubation
b
dsDNA
+ tCTS
Relative Viability (vs no pre-incubation)
Pre-Incubation
No Pre-Incubation
c
On-Target RNP
Off-Target RNP
HDRT Only
Relative Viabilty (vs dsDNA)
Rab11A
CD4
TUBA1B
ACTB
FBL
CLTA
IL2RA
IL2RG
d
NLS
No NLS
Relative Viability (vs dsDNA + Cas9-NLS)
HDRT:
dsDNA
+ tCTS

**Supplementary Fig. 5: Viability loss with Cas9 shuttle requires pre-incubation with HDR template, an on-target gRNA, and partially an NLS peptide.**

(**a**) Pre-incubation of the Cas9 RNP along with dsDNA HDR template containing tCTS on the ends of its homology arms was required for the improvement in knockin efficiency. Pre-incubation was performed for a minimum of 30 seconds at room temperature. In no pre-incubation conditions, the dsDNA HDR template was first mixed with cells, followed by mixing with Cas9 RNP immediately prior to electroporation.

(**b**) Pre-incubation was also necessary for the observed drop in viability when comparing tCTS-modified HDR template to unmodified dsDNA.

(**c**) Use of an on-target gRNA in the Cas9 RNP (specific for both cutting at the desired genomic locus and for binding to the tCTS-binding site introduced on the edges of the HDR template homology arms) was necessary for the observed drop in viability with the Cas9 'shuttle' at certain genomic sites. Note that there was no drop in viability compared to unmodified dsDNA HDR template when electroporating the tCTS-modified HDR template by itself or with an off-target scrambled RNP, indicating the decreased viability was not an inherent property of the tCTS-modified dsDNA template but rather due to the specific complexing with the on-target Cas9 RNP. This reduced viability was not observed at all genomic loci tested.

(**d**) Use of a Cas9 RNP that did not possess an NLS sequence for the 'shuttle' still showed reduced viability when compared to unmodified dsDNA HDR template.

The relative rates of knockin **(a**) or viability (**b-d**) with the various forms of the Cas9 'shuttle' are displayed compared to unmodified dsDNA HDR template (**a-d**) for TCR replacement at the *TRAC* locus (1.5 kb knockin, **a-b, d**) or GFP or tNGFR knockin at the indicated locus (**c**) in multiple technical replicates from n=2 donors (**a-d**). Knockin was measured 4 days post electroporation and viability (total number of live cells relative to no electroporation control) at 2 days post electroporation.

**a**

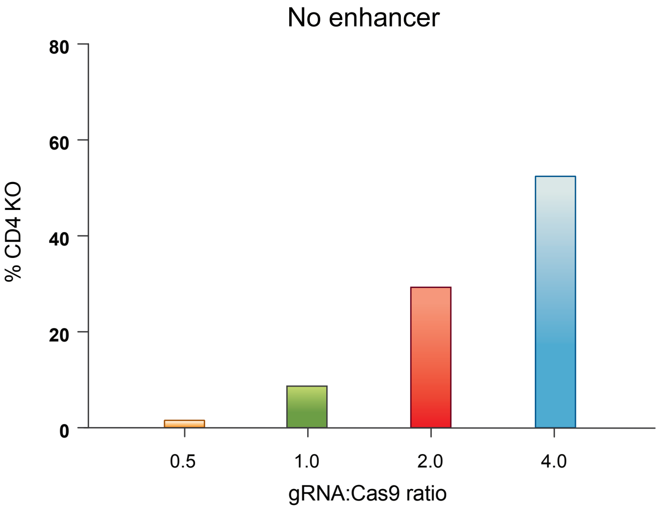


**b**

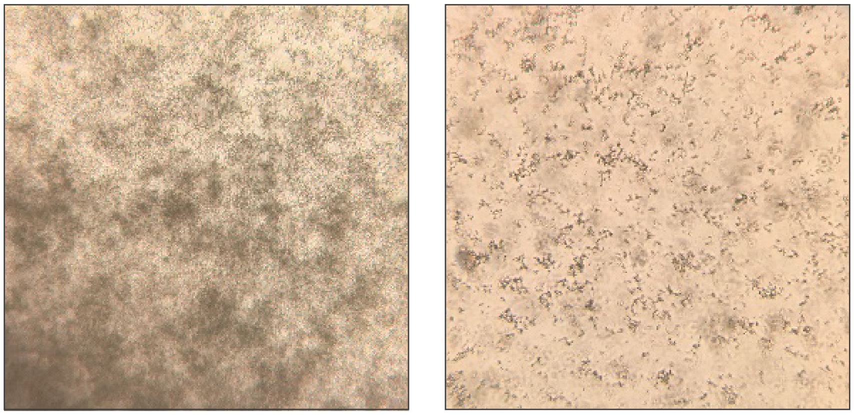

gRNA:Cas9 0.33

gRNA:Cas9 0.33
+ ssODNenh

**c**

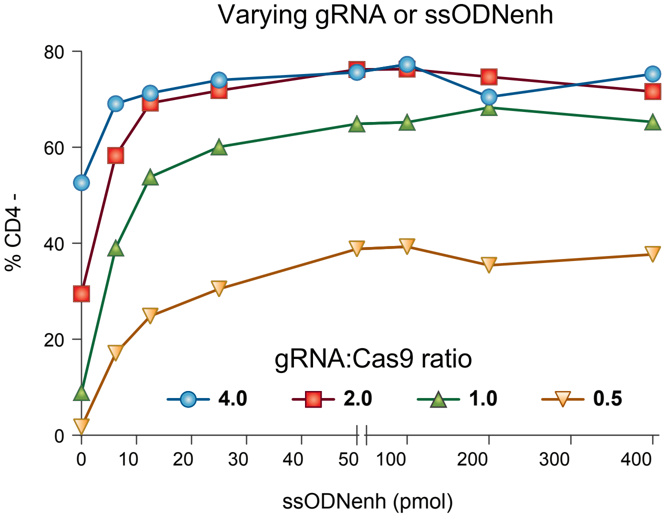

**Supplementary Fig. 6: Increasing gRNA or ssODNenh improved editing efficiency.**

(**a**) Cas9 RNPs targeting *CD4* gene knock-out were electroporated into primary human CD4+ T cells at the indicated gRNA:protein ratio without enhancer or (**c**) with indicated doses of ssODNenh. Loss of surface CD4 expression at 3 days was assessed by flow cytometry. With the exception of 0.5:1 ratio RNPs, which contain only 25pmol of gRNA and thus maximum 50% less functional RNPs; all other gRNA:protein ratios achieved maximal editing with addition of >100pmol of ssODNenh.

(**b**) Photograph of 20x light microscopy showing RNP aggregates at low 0.33:1 ratio RNPs before (left) and within 1 minute (right) after addition of ssODNenh.

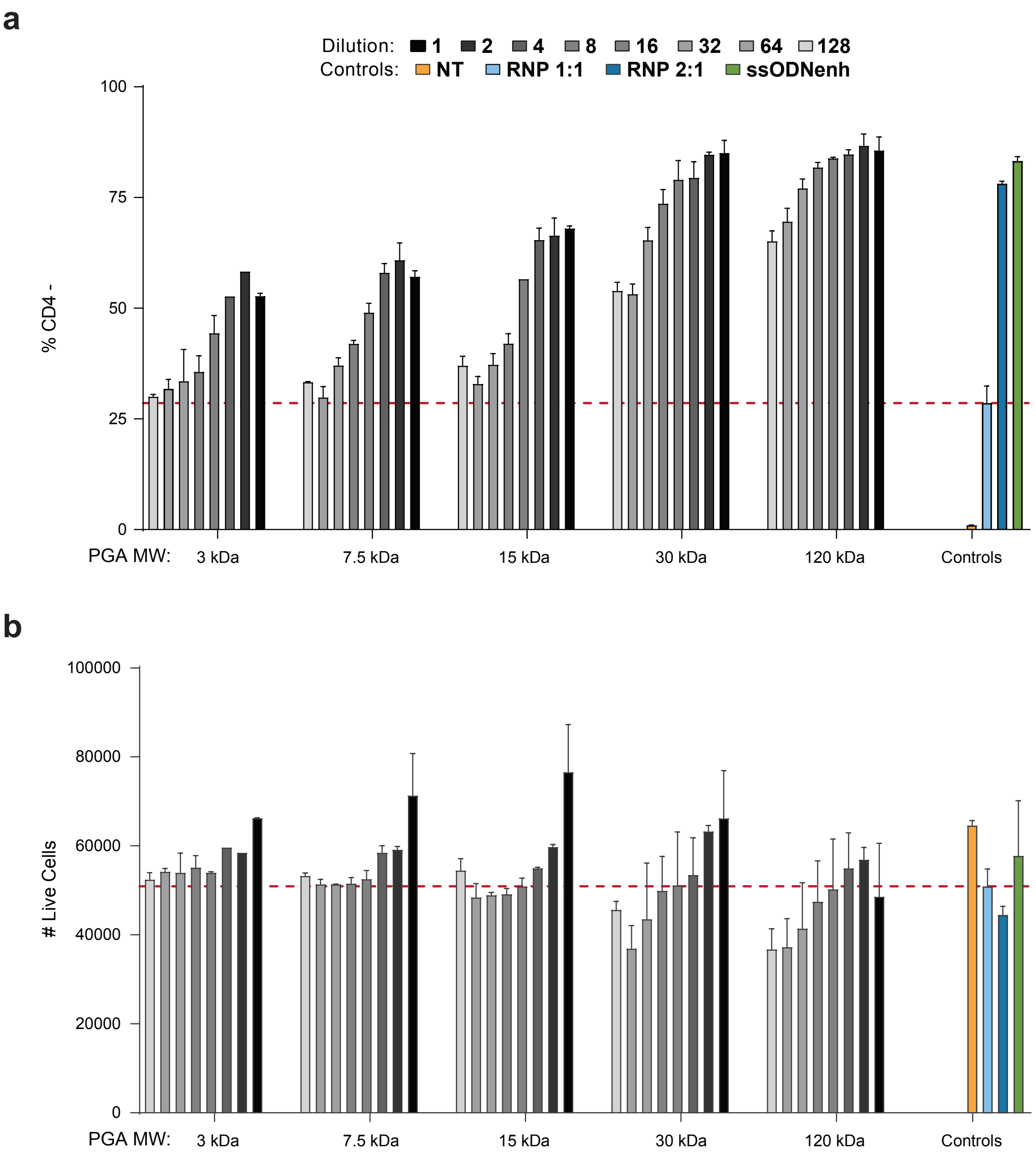
a
Dilution: 1 2 4 8 16 32 64 128
Controls: NT RNP 1:1 RNP 2:1 ssODNenh
% CD4 -
100
75
50
25
0
PGA MW: 3 kDa 7.5 kDa 15 kDa 30 kDa 120 kDa Controls
b
### Live Cells
100000
80000
60000
40000
20000
0
PGA MW: 3 kDa 7.5 kDa 15 kDa 30 kDa 120 kDa Controls

**Supplementary Fig. 7: Poly(glutamic acid) enhanced editing at higher molecular weights**

PGA samples prepared with narrowly-defined molecular weight were assessed for the ability to enhance *CD4* gene knock-out editing. Polymers were reconstituted to 100mg/mL then diluted as indicated, mixed with RNPs formulated at a 1:1 gRNA:protein ratio, and electroporated into primary human CD4+ T cells. **(a)** Loss of surface CD4 expression and **(b)** live cell count at 3 days were assessed by flow cytometry. Higher molecular weight resulted in improved editing but reduced cell viability at the highest MW. Two technical replicates for n=2 different blood donors each averaged with error bars indicating standard deviation.

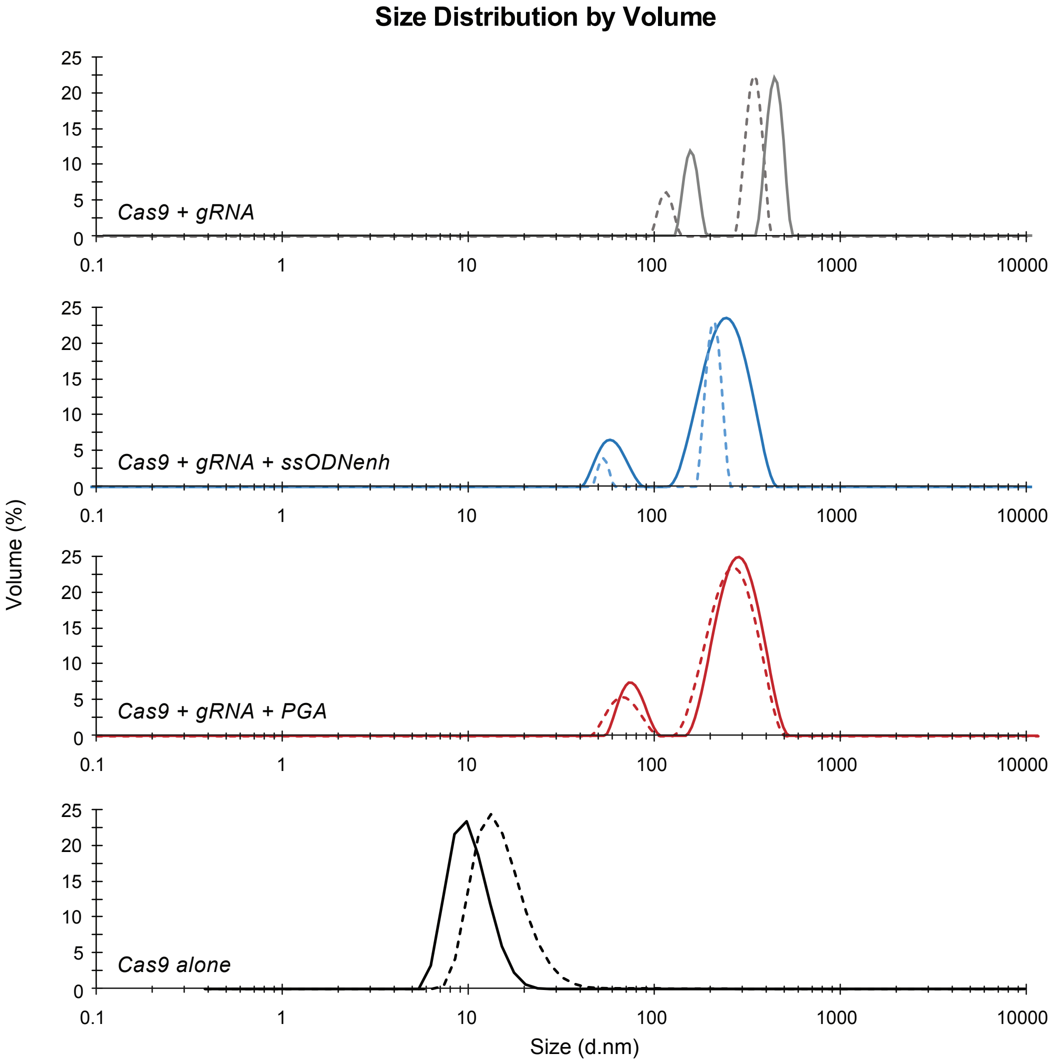


| Sample | Z-avg (nm) | PDI | Peak 1 (nm) | Peak 1 (Volume %) | Peak 2 (nm) | Peak 2 (Volume %) |
|---|---|---|---|---|---|---|
| Cas9 + gRNA 1 | 177.50 | 0.343 | 217.90 | 68.10 | 62.80 | 31.90 |
| Cas9 + gRNA 2 | 195.90 | 0.277 | 173.20 | 80.80 | 46.76 | 19.20 |
| Cas9 + gRNA + ssODNenh 1 | 90.76 | 0.357 | 115.30 | 85.90 | 21.14 | 14.10 |
| Cas9 + gRNA + ssODNenh 2 | 115.00 | 0.183 | 87.88 | 89.50 | 17.04 | 10.50 |
| Cas9 + gRNA + PGA 1 | 77.87 | 0.317 | 116.40 | 87.10 | 23.21 | 12.90 |
| Cas9 + gRNA + PGA 2 | 81.31 | 0.317 | 127.00 | 85.90 | 25.89 | 14.10 |
| Cas9 alone 1 | 25.21 | 0.633 | 10.66 | 99.80 | 309.40 | 0.10 |
| Cas9 alone 2 | 31.83 | 0.44 | 15.80 | 99.50 | 131.00 | 0.50 |

**Supplementary Fig. 8: Anionic polymers stabilized and reduced the hydrodynamic size of RNP nanoparticles.**

Cas9 RNPs were prepared at a 2:1 gRNA:protein ratio, mixed with PGA or ssODNenh, then assessed for particle size by dynamic light scattering. Size distribution by volume % and a summary table of statistics are shown for each of n=2 independent preparations averaged over ten repeated measurements (PDI = polydispersity index).

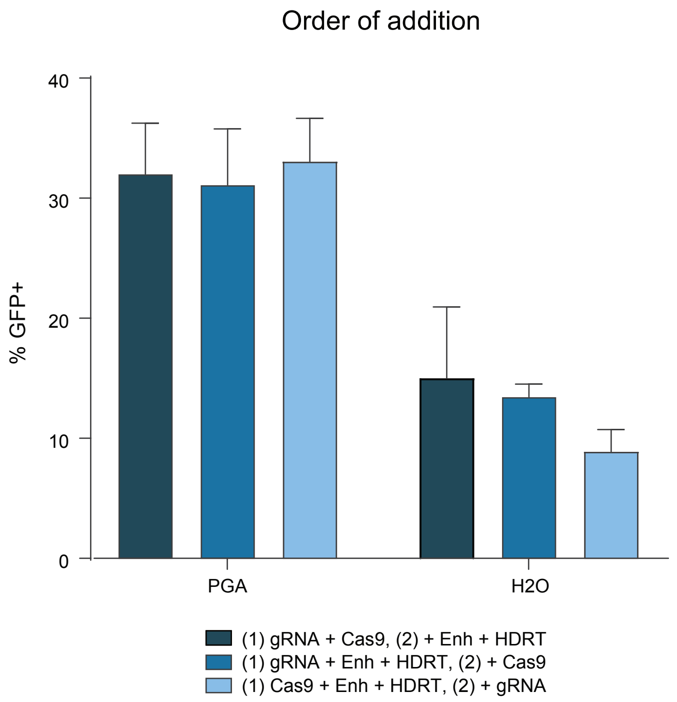

Order of addition
% GFP+
0
10
20
30
40
PGA
H2O
(1) gRNA + Cas9, (2) + Enh + HDRT
(1) gRNA + Enh + HDRT, (2) + Cas9
(1) Cas9 + Enh + HDRT, (2) + gRNA

**Supplementary Fig. 9: Order of addition did not affect editing efficiency.**

Cas9 RNP were formulated at 2:1 gRNA:protein ratio with regular dsDNA HDR template targeting insertion of an N-terminal fusion of GFP to *RAB11A*. PGA polymer or water was added after RNP formation (navy), after gRNA formation but prior to adding Cas9 (blue), or mixed with the protein prior to adding gRNA (light blue). In all three cases, PGA improved editing efficiency but order of addition had no immediately discernable impact. However, we noted improved consistency and workflow when mixing PGA with gRNA first then adding Cas9 protein, and this protocol was adopted for subsequent studies. Two technical replicates for n=2 different blood donors each averaged with error bars indicating standard deviation.

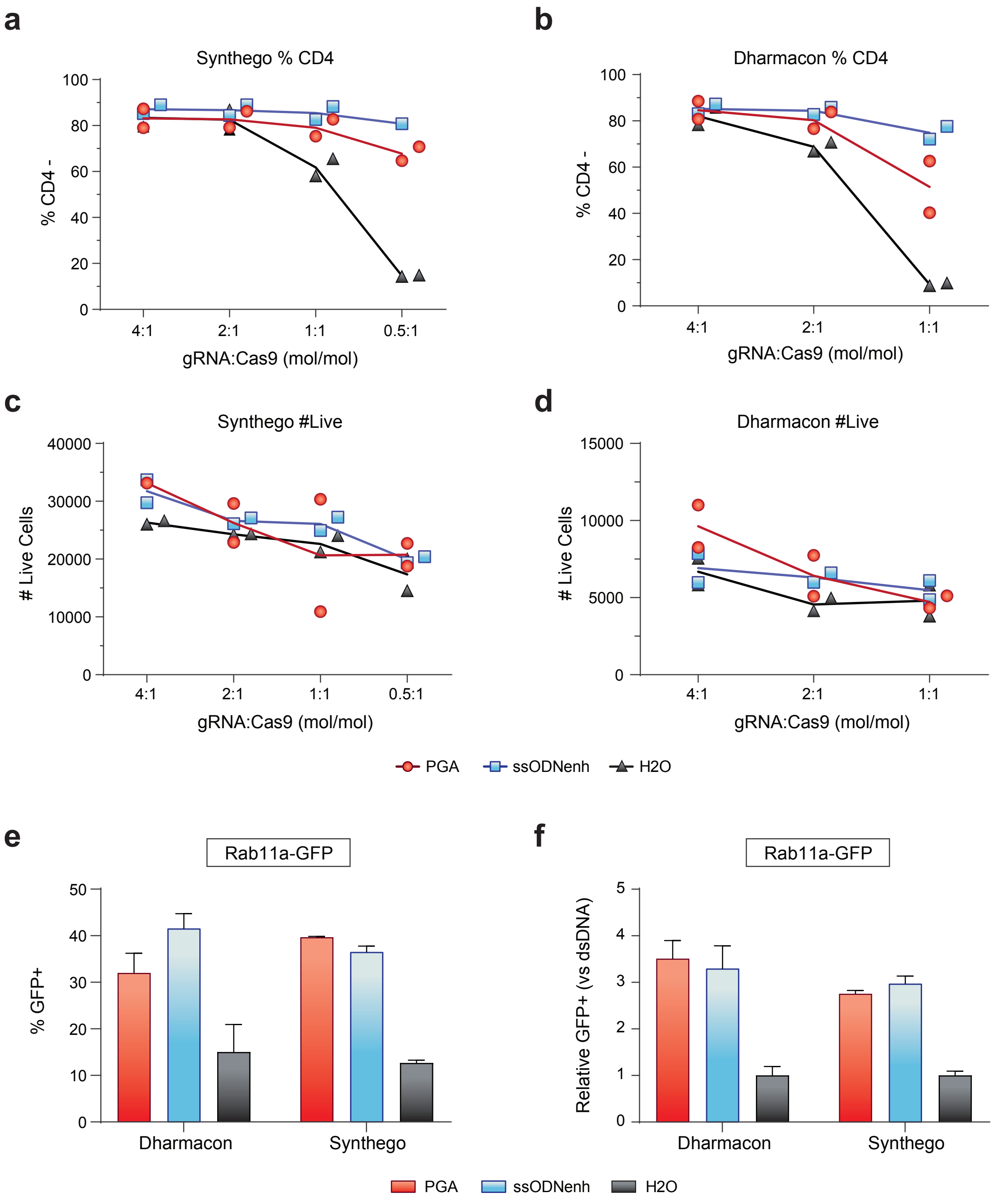
a
Synthego % CD4
% CD4 -
100
80
60
40
20
0
4:1
2:1
1:1
0.5:1
gRNA:Cas9 (mol/mol)
b
Dharmacon % CD4
% CD4 -
100
80
60
40
20
0
4:1
2:1
1:1
gRNA:Cas9 (mol/mol)
c
Synthego #Live
### Live Cells
40000
30000
20000
10000
0
4:1
2:1
1:1
0.5:1
gRNA:Cas9 (mol/mol)
d
Dharmacon #Live
### Live Cells
15000
10000
5000
0
4:1
2:1
1:1
gRNA:Cas9 (mol/mol)
PGA
ssODNenh
H2O
e
Rab11a-GFP
% GFP+
50
40
30
20
10
0
Dharmacon
Synthego
f
Rab11a-GFP
Relative GFP+ (vs dsDNA)
5
4
3
2
1
0
Dharmacon
Synthego
PGA
ssODNenh
H2O

**Supplementary Fig. 10: Both PGA and ssODNenh improved editing independent of guide RNA source.**

Cas9 RNPs were generated at the indicated RNA:protein ratio plus no enhancer (H2O) or with either PGA or ssODNenh and electroporated into primary human CD4+ T cells. RNPs targeted *CD4* gene knock-out (**a-d**), or targeted insertion of an N-terminal fusion of GFP to *RAB11A* when mixed with 1ug of dsDNA HDR template (**e-f**). Loss of surface CD4 expression or GFP positivity and live cell count at 3 days was assessed by flow cytometry. Regardless of guide RNA type either single strand chemically synthesized sgRNA from Synthego (**a,c,e**), or chemically synthesized duplexed tracrRNA:crRNA gRNA from Dharmacon Horizon (**b,d,f**), the addition or either PGA or ssODNenh improved both knock-out and knockin editing efficiency assessed by flow cytometry at day 3. Data shown from each of n=2 different blood donors with line connecting averages (**a-d**); or as the average of n=2 different blood donors with error bars indicating standard deviation (**e-f**).

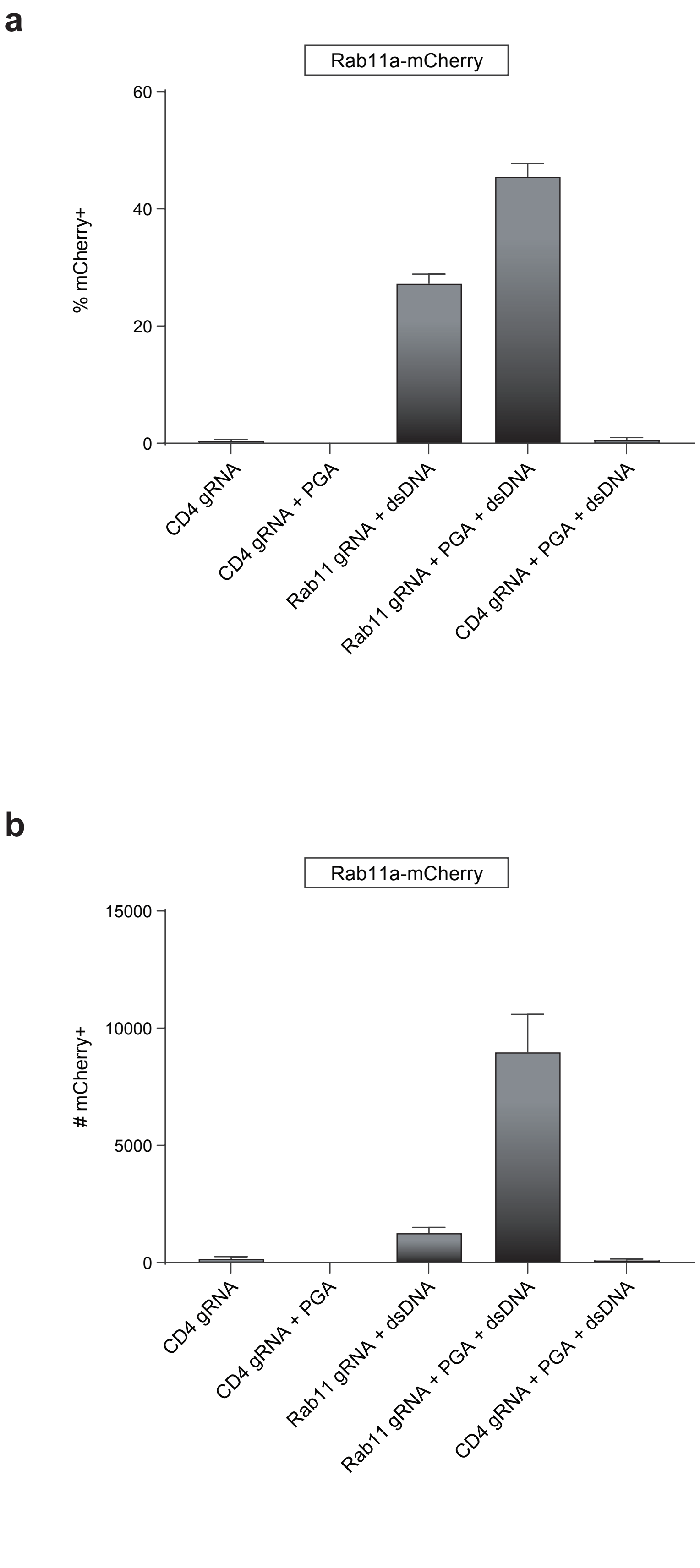
a
Rab11a-mCherry
% mCherry+
60
40
20
0
CD4 gRNA
CD4 gRNA + PGA
Rab11 gRNA + dsDNA
Rab11 gRNA + PGA + dsDNA
CD4 gRNA + PGA + dsDNA
b
Rab11a-mCherry
### mCherry+
15000
10000
5000
0
CD4 gRNA
CD4 gRNA + PGA
Rab11 gRNA + dsDNA
Rab11 gRNA + PGA + dsDNA
CD4 gRNA + PGA + dsDNA

**Supplementary Fig. 11: PGA stabilized HiFi SpyCas9 and improved editing efficiency.**

A novel Cas9 variant, HiFi Cas9, was recently developed with reduced off-target activity but slightly reduced on-target efficiency[14]. HiFiCas9 RNPs were formulated with a guide RNA targeting either the *CD4* gene or *RAB11A* gene with or without PGA polymer at a 2:1 gRNA:protein ratio and electroporated into primary human bulk (CD3+) T cells together with an unmodified dsDNA HDR template for targeted integration of an N-terminal fusion of mCherry to *RAB11A*. (**a**) mCherry-expressing cells and (**b**) live cell count were quantified at day 3 by flow cytometry. PGA improved on-target editing efficiency while maintaining minimal detection of mCherry+ cells when using an off-target gRNA. Data shown as the average of n=2 different blood donors with error bars indicating standard deviation.

**a**

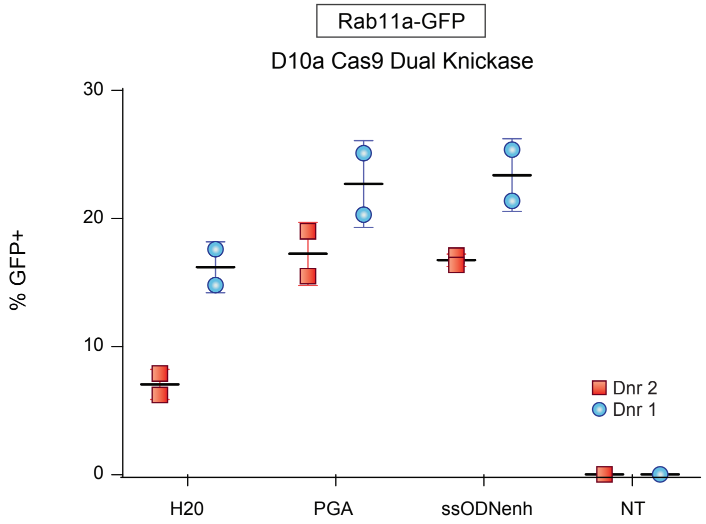

Rab11a-GFP
D10a Cas9 Dual Knickase
30
20
10
0
% GFP+
H20
PGA
ssODNenh
NT
Dnr 2
Dnr 1


**b**

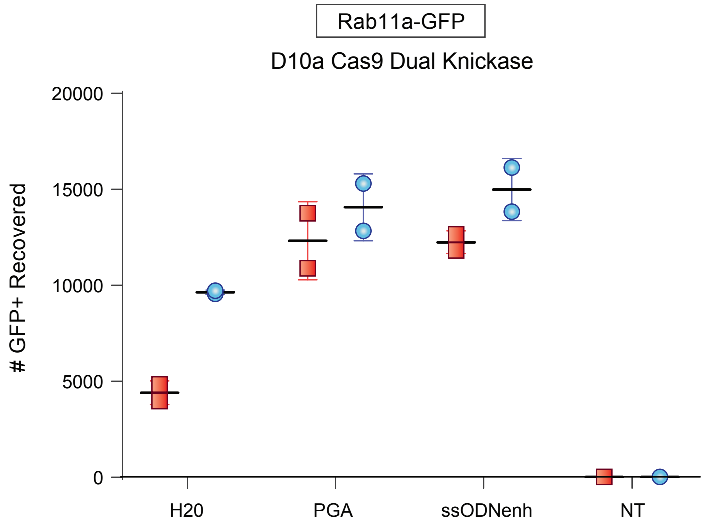

Rab11a-GFP
D10a Cas9 Dual Knickase
20000
15000
10000
5000
0
### GFP+ Recovered
H20
PGA
ssODNenh
NT

**Supplementary Fig. 12: PGA or ssODNenh improved editing efficiency with D10a 'nickase' variant Cas9 RNPs.**

A dual nickase approach to induce neighboring opposed single strand breaks has been proposed as a strategy to minimize off-target editing[15]. D10a Cas9 RNPs were formulated with guide RNAs as previously described[5] targeting the *RAB11A* gene exon 1 with or without polymer at a 2:1 gRNA:protein ratio, and 50 pmol RNP were electroporated into primary human bulk (CD3+) T cells together with 1ug of regular dsDNA HDR template encoding insertion of an N-terminal fusion of GFP to *RAB11A*. (**a**) GFP-expressing cells and (**b**) live cell count were quantified at day 3 by flow cytometry. Two technical replicates each shown for two different blood donors.

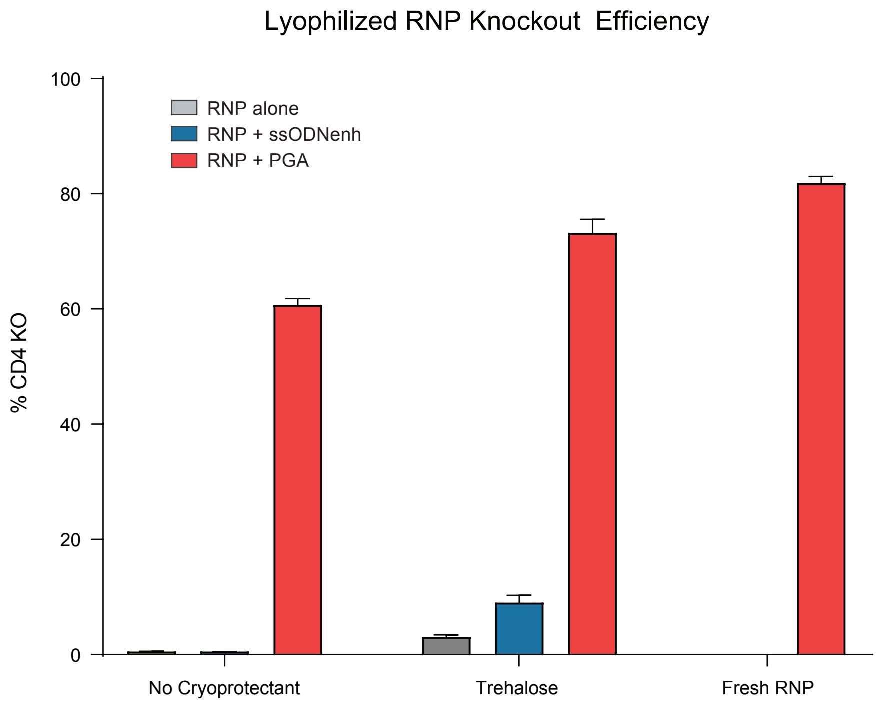

Lyophilized RNP Knockout Efficiency
100
80
60
40
20
0
% CD4 KO
RNP alone
RNP + ssODNenh
RNP + PGA
No Cryoprotectant
Trehalose
Fresh RNP

**Supplementary Fig. 13: PGA-stabilized Cas9 RNP nanoparticles could be lyophilized with retained editing efficiency.**

Cas9 RNPs targeting the *CD4* gene were prepared at 2:1 ratio gRNA:protein without or with PGA or ssODNenh, diluted in water or equal volumes of 50mM trehalose, frozen in liquid nitrogen, lyophilized overnight, then stored dry at -80C. RNPs were later reconstituted in water prior to electroporating into primary human CD4+ T cells. Cas9 RNP nanoparticles stabilized with PGA, but not ssODNenh, were protected through lyophilization and reconstitution and retained robust knock-out editing efficiency compared to freshly prepared RNPs. Data shown as the average of three technical replicates for one blood donor; error bars indicate standard deviation.

**a**

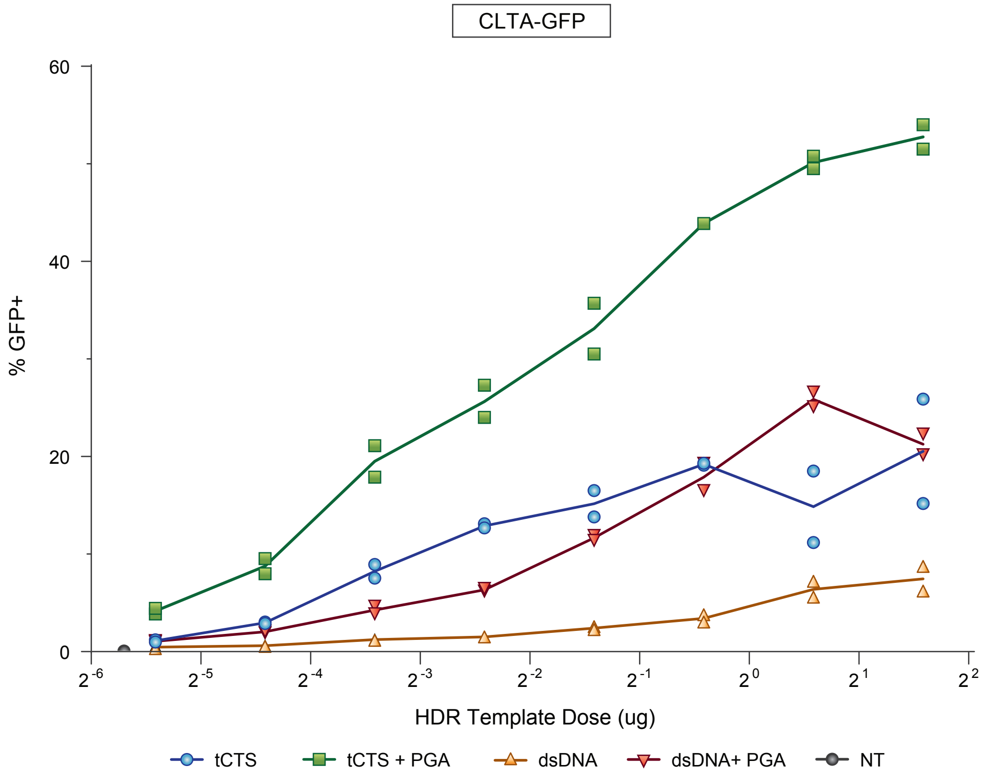
CLTA-GFP
60
40
20
0
% GFP+
HDR Template Dose (ug)
tCTS
tCTS + PGA
dsDNA
dsDNA+ PGA
NT

**b**

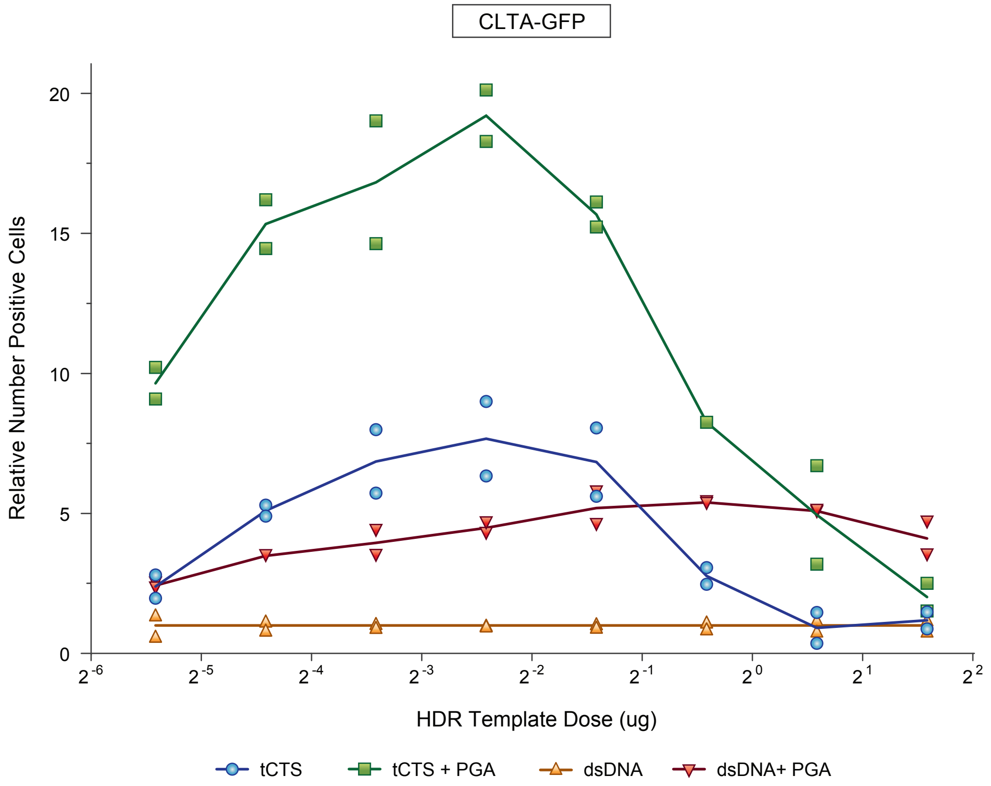
CLTA-GFP
20
15
10
5
0
Relative Number Positive Cells
HDR Template Dose (ug)
tCTS
tCTS + PGA
dsDNA
dsDNA+ PGA

**Supplementary Fig. 14: PGA-stabilized Cas9 RNP nanoparticles improved knockin.**

A series of doses of unmodified HDR template or tCTS-modified HDR template targeting a N-terminal fusion of GFP to Clathrin were combined with 50 pmol of regular Cas9 RNP or PGA-stabilized Cas9 RNP then electroporated in primary human bulk (CD3+) T cells. (**a**) GFP-expressing cells and (**b**) quantity recovered GFP+ cells were quantified at day 3 by flow cytometry. Data shown from two different blood donors with lines connecting means for each condition.

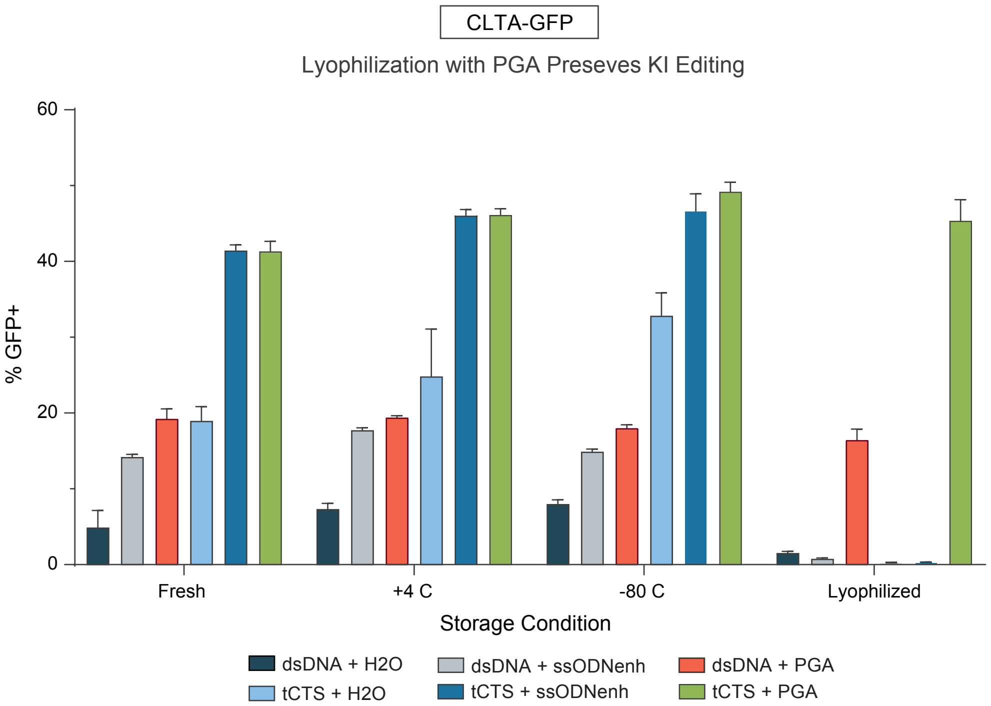

CLTA-GFP
Lyophilization with PGA Preseves KI Editing
60
40
20
0
% GFP+
Fresh
+4 C
-80 C
Lyophilized
Storage Condition
dsDNA + H2O
dsDNA + ssODNenh
dsDNA + PGA
tCTS + H2O
tCTS + ssODNenh
tCTS + PGA

**Supplementary Fig. 14: PGA-stabilized Cas9 RNP nanoparticles complexed with tCTS-modified 'shuttle' HDR template could be lyophilized with retained editing efficiency.**

Cas9 RNPs were prepared at 2:1 ratio gRNA:protein together with extra H2O, PGA, or ssODNenh, then mixed with regular dsDNA HDR template (ds) or tCTS-modified HDR template (tCTS) targeting insertion of an N-terminal fusion of GFP to *RAB11A*. Samples were incubated at 37C to enhance RNP-HDR template interaction, diluted in equal volumes of 50mM trehalose, frozen in liquid nitrogen, lyophilized overnight, then stored at -80C. RNP-HDR templates were later reconstituted in water prior to electroporating into primary human bulk (CD3+) T cells. Cas9 RNP-HDR template nanoparticles stabilized with PGA, but not ssODNenh, were protected through lyophilization and reconstitution and retained robust knock-out editing efficiency compared to freshly prepared RNPs. Data shown as the average of three technical replicates for one blood donor; error bars indicate standard deviation.

**Supplementary Table 1**.

List of HDR Template, DNA primer, and gRNA sequences used in study. For sequences please see uploaded MS Excel file.

**Supplementary Table 2 - Polymers Tested**

| Polymer | Abbreviation | MW (kDa) | Concentration (mg/mL) | Manufacturer |
|---|---|---|---|---|
| polyethylenimine (branched) | bPEI | 2 | 5 | Sigma |
| poly(L-arginine) HCL | PLA | 15-70 | 100 | Sigma |
| poly(L-lysine) HBr | PLL | 15-30 | 100 | Sigma |
| poly(L-ornithine) HBr | PLO | 30-70 | 100 | Sigma |
| Protamine Sulfate | PS | >100 U/mg | 10 | Fresenius Kabi |
| Poly(ethylene glycol) | PEG-35 | 35 | 44 | Sigma |
| Poly(L-glutamic acid) | PGA | 15-50 | 100 | Sigma |
| Poly(L-glutamic acid) | PLE20 | 3 | 100 | Alamanda Polymers |
| Poly(L-glutamic acid) | PLE50 | 7.5 | 100 | Alamanda Polymers |
| Poly(L-glutamic acid) | PLE100 | 15 | 100 | Alamanda Polymers |
| Poly(L-glutamic acid) | PLE200 | 30 | 100 | Alamanda Polymers |
| Poly(L-glutamic acid) | PLE800 | 120 | 100 | Alamanda Polymers |
| Heparin | Hep | >180 U/mg | 60 | Sigma |
| Hyaluronic acid | HA-150 | 100-150 | 75 | Lifecore Biomedical |
| Poly(acrylic acid) | PAA-5 | 5 | 180 | Sigma |
| Poly(acrylic acid) | PAA-25 | 25 | 100 | Sigma |
| Poly(acrylic acid) | PAA-250 | 250 | 72 | Sigma |
| poly(L-aspartic acid) | PLD | 27 | 13.3 | Alamanda Polymers |
| ssODNenh electroporation enhancer* | ssODNenh | 31 | 3.1 | IDT Technologies |

* ssODNenh sequence:

TTAGCTCTGTTTACGTCCCAGCGGGCATGAGAGTAACAAGAGGGTGTGGTAATATTACGG
TACCGAGCACTATCGATACAATATGTGTCATACGGACACG

Positively-charged polymers shaded in yellow, neutral polymers shaded in green, negatively-charged polymers shaded in blue.

**Supplementary Table 3 – Antibodies Used in Study**

| Target | Fluor | Vendor | Clone | Catalog Number |
|---|---|---|---|---|
| Fc Receptor | - | Biolegend | Human TruStain FcX | 422302 |
| GhostDye Red 780 | - | Tonbo | - | 13-0865-T500 |
| GhostDye Violet 510 | - | Tonbo | - | 13-0870-T500 |
| LIVE/DEAD® Fixable Aqua Dead Cell Stain | - | ThermoFisher Scientific | - | L34966 |
| CD19 | PacBlue | Biolegend | HIB19 | 302223 |
| CD271 (tNGFR) | APC | Biolegend | ME20.4 | 345108 |
| CD3 | AlexaFluor 700 | Becton-Dickson | UCHT1 | 557943 |
| CD3 | PE | Biolegend | UCHT1 | 300408 |
| CD34 | PE-Cy7 | Becton-Dickson | 8G12 | 348791 |
| CD4 | PE-Cy7 | Biolegend | OKT4 | 317414 |
| CD4 | FITC | Biolegend | SK3 | 344604 |
| CD4 | PerCP | Tonbo | SK3 | 67-0047-T500 |
| CD56 | PerCP | Biolegend | HCD56 | 318342 |
| CD8 | APC | Tonbo | OKT8 | 20-0086-T100 |
| CD8 | PE-Cy7 | Becton-Dickson | SK1 | 335787 |
| TCR- 1G4 | PE | Immudex | HLA-A*0201/SLLMWITQV | WB3247-PE |
| TCR-gd | PE-Cy7 | Biolegend | B1 | 331221 |
| TCR-αβ | BV-421 | Biolegend | IP26 | 306722 |
